# The Maximum-Tolerated Dose and Pharmacokinetics of ISFP10 a Novel Inhibitor of Fungal Phosphoglucomutase (PGM)

**DOI:** 10.1101/2025.06.17.660114

**Authors:** Theodore J. Kottom, Andrew H. Limper

**Author notes:** Address correspondence to Dr. Theodore Kottom, 8-23 Stabile, Mayo Clinic, Rochester, MN 55905.

## Abstract

**Background:** The newly identified small molecule ISFP10 has demonstrated the ability to inhibit fungal phosphoglucomutases (PGM), leading to decreased fungal growth and survival. Exhibiting 50 times greater selectivity for fungal PGM compared to the human homolog, ISFP10 shows promise as a broad-spectrum antifungal and a candidate for preliminary testing in rodent models of fungal infections, including Pneumocystis pneumonia (PCP). In preparation for targeted and relevant administration of the drug in fungal mouse models of infection, the current study describes the maximum-tolerated-dose (MTD) and pharmacokinetics (PK) of ISFP10 in mice.

**Methods:** For the MTD study, 24 C57BL/6 mice were randomly distributed into 6 groups (2M/2F per group): Placebo (vehicle; 10% DMSO with 90% of 0.5% methylcellulose in 0.9% NaCl) at various doses (12.5, 25, 50, 100, 200 mg/kg), were administered twice daily by intraperitoneally (IP) for 7 consecutive days. For the PK study, 108 mice were administered either vehicle or ISFP10 in single-dose intraperitoneal (IP) injections twice daily for 7 consecutive days at concentrations of 0, 12.5, 25, 50, and 100 mg/kg. Pharmacokinetics were assessed in plasma, and epithelial lining fluid (ELF). Analysis of this test compound was performed using LC-MS/MS for PK evaluation on blood samples drawn from mice at multiple time points.

**Results:** IP administered, ISFP10 did not produce significant changes in body weight, food consumption or adverse events in the MTD up to 100 mg/kg dosing. The plasma PK study demonstrated T_max_ values ranging from 2 to 8 h. Non-compartmental analysis (NCA) yielded elimination half-life values, t_1/2_, of 4.39 and 5.31 h for the 12.5 and 50 mg/kg dose groups. Concentration dependent peak-blood concentration (C_max_) at the dosing and length tested ranged from 3.450-5.140 ug/ml.

**Conclusions:** In conclusion, ISFP10, administered intraperitoneally to mice twice daily at doses up to 100 mg/kg, exhibited no inherent safety concerns based on the analyzed parameters. Pharmacokinetic analysis revealed slow absorption and distribution kinetics, with plasma T_max_ values ranging from 2 to 8 hours and elimination half-lives of approximately 4–5 hours at lower doses. These data support broader *in vivo* testing of the inhibitor as potential novel antifungal in fungal diseases including PCP.

## 1. Introduction

The fungal cell wall is an attractive target for antifungal therapies because it has no equivalent structure in human cells. Among its key components, β-glucans are particularly abundant and play a crucial role in maintaining cell wall rigidity. The primary forms present are β-1,3-and β-1,6-glucans [1], which are synthesized from UDP-glucose by their respective synthases [1, 2]

In carbohydrate metabolism, phosphoglucomutase (PGM) is essential for converting glucose-1-phosphate to glucose-6-phosphate, an important step in the synthesis of UDP-glucose, the precursor to fungal cell wall β-glucan synthesis, and the formation of other carbohydrates [3, 4]. PGM has been identified as a potential therapeutic target in *Aspergillus fumigatus*, with inhibition demonstrated using a para-aryl derivative known as ISFP10 [4]. Our laboratory has further shown that PGM is also a viable therapeutic target in *Pneumocystis jirovecii* and *Pneumocystis murina* [3, 5], providing preliminary evidence that ISFP10 may serve as an effective antifungal agent.

In order to test ISFP10 in clinically relevant animal fungal models of infection, data on its maximum-tolerated dose (MTD) and pharmacokinetics (PK) in animals must be established. MTD and PK data of ISFP10 following intraperitoneally (IP) administration of various doses ranging from 12.5 mg/kg twice daily to 200 mg/kg twice daily was conducted in male and female C57BL/6 mice. Results from both these studies demonstrated that ISFP10 can be administered at doses as high as 100mg/kg 2X/day in mice. This study also describes the pharmacokinetics of ISFP10 and provides initial important information of its use a novel antifungal in the treatment of animal models of fungal infection.

## 2. Methods

### 2.1. Animals

Male and female immunocompetent C57BL/6 mice at 7-8 weeks of age were used for all experiments.

### 2.2. Administration of ISFP10

#### MTD Analysis

The MTD study was performed according to previously published methods [6]. ISFP10 and the vehicle control were administered intraperitoneally (IP) to animals twice daily (BID) for 7 consecutive days at 12-hour intervals at 12.5, 25, 50, 100, and 200 mg/kg. The dosing formulations were prepared using 10% DMSO and 90% of 0.5% methyl cellulose (MC) in 0.9% NaCl. All formulations were visually inspected for homogeneity and were consistently observed as white emulsions. Animals were monitored 30 minutes post-dosing for signs of acute toxicity, which were recorded if present. In cases of severe acute toxicity, animals were humanely euthanized.

#### Clinical Examinations

Within 30 minutes following IP administration, animals were examined for acute toxic effects—including mortality, convulsions, tremors, muscle relaxation, and sedation—as well as autonomic responses such as diarrhea, salivation, lacrimation, vasodilation, and piloerection. Additional cage-side observations included assessments of gait, posture, coat condition (e.g., ruffled fur), activity level (e.g., immobility), mucous membrane status, salivation, tremors, convulsions, response to handling, respiratory function, stool output, mortality, and body weight. These parameters were recorded both after each subsequent IP dose and daily for three days following the final administration. Animals exhibiting signs of distress or moribundity were euthanized humanely at earlier time points as necessary.

#### PK Analysis

PK analysis in mice was conducted similarly to previously published methods [7–9]. The ISFP10 dose solution was freshly prepared by first dissolving the compound in 100% DMSO, followed by addition of 0.5% methyl cellulose (MC) in 0.9% NaCl at a 10% to 90% vol/vol ratio to make the final DMSO concentration of 10% and methyl cellulose concentration of 0.45% in 0.9% NaCl. All formulations were examined visually for uniformity and consistently appeared as white emulsions. ISFP10 and vehicle control were administered intraperitoneally (IP) to animals twice daily (BID) for 7 consecutive days at 12-hour intervals at 12.5, 25, 50, and 100 mg/kg. Analysis of this test compound was performed using LC-MS/MS for PK evaluation on blood samples drawn from mice at multiple time points.

### 2.3. Study results and discussion

#### Tolerability assessment of ISFP10 in immunocompetent C57BL/6 mice

The tolerability of ISFP10 was evaluated in immunocompetent, uninfected male and female C57BL/6 mice. The compound was administered intraperitoneally (IP) twice daily at 12-hour intervals (BID, q12h) for seven consecutive days at escalating dose levels of 12.5, 25, 50, 100, and 200 mg/kg, as outlined in **Table 1**. Following the initial dose, animals were monitored for signs of acute toxicity (**Table 2**). Clinical symptoms were assessed and scored after each administration and continued daily for three days following the final dose, with results summarized in **Table 3 (1-18**). Body weight was recorded daily and is presented in **Table 4 (1-2**). Mortality outcomes across dose groups are reported in **Table 5 (1-2**).

**Table 1.** Tolerability analysis, uninfected C57BL/6 mice, study design:

| Group | Test Substance | Dose Schedule | Dose Route | Dosage |  |  |  | Mice |  |
| --- | --- | --- | --- | --- | --- | --- | --- | --- | --- |
|  |  |  |  | Conc. (mg/mL) | Dose Volume (mL/kg) | Dose (mg/kg) | Dose/day (mg/kg/24 h) | Male | Female |
| 1 | Vehicle | BID q12h x 7 days | IP | -- | 10 | -- | -- | 2 | 2 |
| 2 | ISFP10 | BID q12h x 7 days | IP | 1.25 | 10 | 12.5 | 25 | 2 | 2 |
| 3 | ISFP10 | BID q12h x 7 days | IP | 2.5 | 10 | 25 | 50 | 2 | 2 |
| 4 | ISFP10 | BID q12h x 7 days | IP | 5 | 10 | 50 | 100 | 2 | 2 |
| 5 | ISFP10 | BID q12h x 7 days | IP | 10 | 10 | 100 | 200 | 2 | 2 |
| 6 | ISFP10 | BID q12h x 7 days | IP | 20 | 10 | 200 | 400 | 2 | 2 |
| C57BL/6 mice: |  |  |  |  |  |  |  | 24 |  |
| Vehicle was 10% DMSO with 90% of 0.5% MC in 0.9% NaCl. |  |  |  |  |  |  |  |  |  |
| The animals were intraperitoneally (IP) administered twice at 12 h intervals (BID q12h) for 7 consecutive days (x 7 days). |  |  |  |  |  |  |  |  |  |
| Full clinical examinations (Table 2) were conducted at 30 minutes after the administration. Cage side observations were conducted at 30 min after each subsequent administration as well as daily for 3 days after the last administration. A final observation of mortality and body weight was conducted and recorded 3 days following the last dose administration. |  |  |  |  |  |  |  |  |  |
| The six groups were conducted concurrently. |  |  |  |  |  |  |  |  |  |

**Table 2.** Clinical symptoms for observation:

| Body Weight (B.W.) | Decrease in Touch Response | Decrease in Spontaneous Activity | Low Limb Post | Body Temperature |
| --- | --- | --- | --- | --- |
| Irritability | Increase in Exploration | Straub Tail | Skin Color | Piloerection |
| Hyperactivity | Decrease in Exploration | Reactivity | Respiration | Increase in Palpebral Size |
| Increase in Startle Response | Pinna | Righting | Salivation (Fluid and Viscosity) | Decrease in Palpebral Size |
| Increase Touch Response | Placing | Ataxia | Lacrimation | Death |
| Decrease Startle Response | Tremor | Convulsion (Chronic / Tonic) | Diarrhea |  |
| Clinical observations were made at 30 minutes after the IP dose administration and recorded. |  |  |  |  |

**Table 3-1.** MTD results of ISFP10, clinical examination at 30 minutes after the 1^st^ IP dosing, female.

| Test Article | Vehicle |  | PT# 1288839<br>(NID-490)<br>(ISFP10) |  |  |  |  |  |  |  |  |  |
| --- | --- | --- | --- | --- | --- | --- | --- | --- | --- | --- | --- | --- |
| Route/Schedule | IP, BID, q12h<br>x 7 days |  | IP, BID, q12h<br>x 7 days |  |  |  |  |  |  |  |  |  |
| Dosage | 10 mL/kg |  | 12.5 mg/kg |  | 25 mg/kg |  | 50 mg/kg |  | 100 mg/kg |  | 200 mg/kg |  |
| BEHAVIORAL | #1 | #2 | #1 | #2 | #1 | #2 | #1 | #2 | #1 | #2 | #1 | #2 |
| Body Weight (g) | 20 | 18 | 19 | 20 | 21 | 19 | 20 | 20 | 19 | 20 | 21 | 21 |
| Irritability | - | - | - | - | - | - | - | - | - | - | - | - |
| Hyperactivity | - | - | - | - | - | - | - | - | - | - | - | - |
| Inc. Startle | - | - | - | - | - | - | - | - | - | - | - | - |
| Inc. Touch | - | - | - | - | - | - | - | - | - | - | - | - |
| Dec. Startle Response | - | - | - | - | - | - | - | - | - | - | - | - |
| Dec. Touch Response | - | - | - | - | - | - | - | - | - | - | - | - |
| Inc. Exploration | - | - | - | - | - | - | - | - | - | - | - | - |
| Dec. Exploration | - | - | - | - | - | - | - | - | - | - | - | - |
| Pinna | - | - | - | - | - | - | - | - | - | - | - | - |
| Placing | - | - | - | - | - | - | - | - | - | - | - | - |
| NEUROLOGIC |  |  |  |  |  |  |  |  |  |  |  |  |
| Tremor | - | - | - | - | - | - | - | - | - | - | - | - |
| Dec. Spont. Activity | - | - | - | - | - | - | - | - | - | - | - | - |
| Straub Tail | - | - | - | - | - | - | - | - | - | - | - | - |
| Reactivity | - | - | - | - | - | - | - | - | - | - | - | - |
| Righting | - | - | - | - | - | - | - | - | - | - | - | - |
| Ataxia | - | - | - | - | - | - | - | - | - | - | - | - |
| Convulsion C.T.C-T | - | - | - | - | - | - | - | - | - | - | - | - |
| Low Limb Post | - | - | - | - | - | - | - | - | - | - | - | - |
| Abdominal Tone | - | - | - | - | - | - | - | - | ± | ± | ± | ± |
| Limb Tone | - | - | - | - | - | - | - | - | - | - | - | - |
| Grip Strength | - | - | - | - | - | - | - | - | - | - | - | - |
| AUTONOMIC |  |  |  |  |  |  |  |  |  |  |  |  |
| Skin Color | - | - | - | - | - | - | - | - | - | - | - | - |
| Respiration | - | - | - | - | - | - | - | - | - | - | - | - |
| Salivation F.V. | - | - | - | - | - | - | - | - | - | - | - | - |
| Lacrimation | - | - | - | - | - | - | - | - | - | - | - | - |
| Diarrhea | - | - | - | - | - | - | - | - | - | - | - | - |
| Body Temperature | - | - | - | - | - | - | - | - | - | - | - | - |
| Piloerection | - | - | - | - | - | - | - | - | - | - | - | - |
| Inc. Palpebral Size | - | - | - | - | - | - | - | - | - | - | - | - |
| Dec. Palpebral Size | - | - | - | - | - | - | - | - | - | - | - | - |
| Others | - | - | - | - | - | - | - | - | - | - | - | - |
| Death | - | - | - | - | - | - | - | - | - | - | - | - |

**Notes (alphabetical order):**

**Table 3-2.** MTD results of ISFP10, clinical examination at 30 minutes after the 1^st^ IP dosing, male.

| Test Article | Vehicle |  | PT# 1288839<br>(NID-490)<br>(ISFP10) |  |  |  |  |  |  |  |  |  |
| --- | --- | --- | --- | --- | --- | --- | --- | --- | --- | --- | --- | --- |
| Route/Schedule | IP, BID, q12h<br>x 7 days |  | IP, BID, q12h<br>x 7 days |  |  |  |  |  |  |  |  |  |
| Dosage | 10 mL/kg |  | 12.5 mg/kg |  | 25 mg/kg |  | 50 mg/kg |  | 100 mg/kg |  | 200 mg/kg |  |
| BEHAVIORAL | #1 | #2 | #1 | #2 | #1 | #2 | #1 | #2 | #1 | #2 | #1 | #2 |
| Body Weight (g) | 23 | 24 | 24 | 23 | 23 | 22 | 23 | 24 | 20 | 22 | 22 | 21 |
| Irritability | - | - | - | - | - | - | - | - | - | - | - | - |
| Hyperactivity | - | - | - | - | - | - | - | - | - | - | - | - |
| Inc. Startle | - | - | - | - | - | - | - | - | - | - | - | - |
| Inc. Touch | - | - | - | - | - | - | - | - | - | - | - | - |
| Dec. Startle Response | - | - | - | - | - | - | - | - | - | - | - | - |
| Dec. Touch Response | - | - | - | - | - | - | - | - | - | - | - | - |
| Inc. Exploration | - | - | - | - | - | - | - | - | - | - | - | - |
| Dec. Exploration | - | - | - | - | - | - | - | - | - | - | - | - |
| Pinna | - | - | - | - | - | - | - | - | - | - | - | - |
| Placing | - | - | - | - | - | - | - | - | - | - | - | - |
| NEUROLOGIC |  |  |  |  |  |  |  |  |  |  |  |  |
| Tremor | - | - | - | - | - | - | - | - | - | - | - | - |
| Dec. Spont. Activity | - | - | - | - | - | - | - | - | - | - | - | - |
| Straub Tail | - | - | - | - | - | - | - | - | - | - | - | - |
| Reactivity | - | - | - | - | - | - | - | - | - | - | - | - |
| Righting | - | - | - | - | - | - | - | - | - | - | - | - |
| Ataxia | - | - | - | - | - | - | - | - | - | - | - | - |
| Convulsion C.T.C-T | - | - | - | - | - | - | - | - | - | - | - | - |
| Low Limb Post | - | - | - | - | - | - | - | - | - | - | - | - |
| Abdominal Tone | - | - | - | - | - | - | - | - | ± | ± | ± | ± |
| Limb Tone | - | - | - | - | - | - | - | - | ± | ± | ± | ± |
| Grip Strength | - | - | - | - | - | - | - | - | - | - | - | - |
| AUTONOMIC |  |  |  |  |  |  |  |  |  |  |  |  |
| Skin Color | - | - | - | - | - | - | - | - | - | - | - | - |
| Respiration | - | - | - | - | - | - | - | - | - | - | - | - |
| Salivation F.V. | - | - | - | - | - | - | - | - | - | - | - | - |
| Lacrimation | - | - | - | - | - | - | - | - | - | - | - | - |
| Diarrhea | - | - | - | - | - | - | - | - | - | - | - | - |
| Body Temperature | - | - | - | - | - | - | - | - | - | - | - | - |
| Piloerection | - | - | - | - | - | - | - | - | - | - | - | - |
| Inc. Palpebral Size | - | - | - | - | - | - | - | - | - | - | - | - |
| Dec. Palpebral Size | - | - | - | - | - | - | - | - | - | - | - | - |
| Others | - | - | - | - | - | - | - | - | - | - | - | - |
| Death | - | - | - | - | - | - | - | - | - | - | - | - |

**Notes (alphabetical order):**

**Table 3-3.**
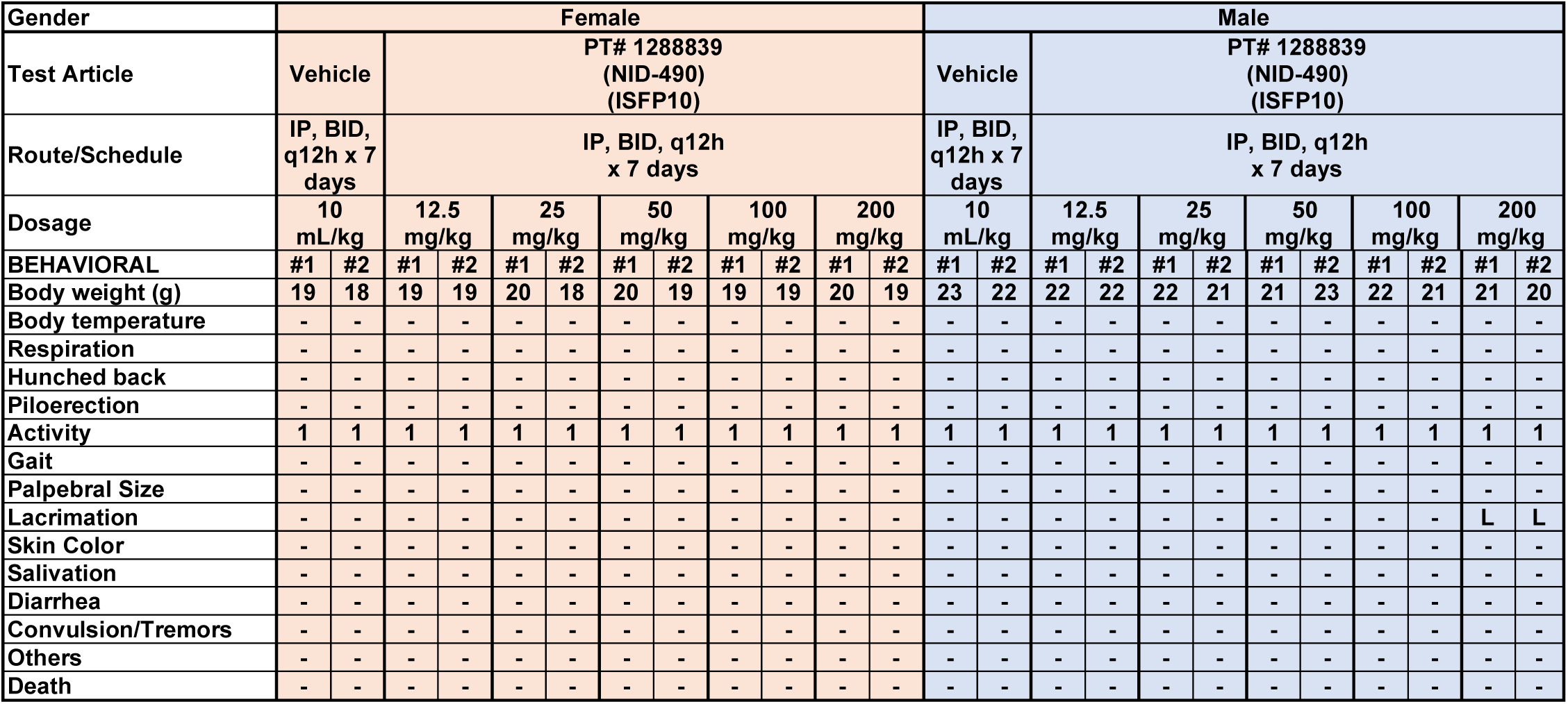
MTD results of ISFP10, cage side observation at 30 minutes after the 2^nd^ IP dosing.

**Table 3-4.**
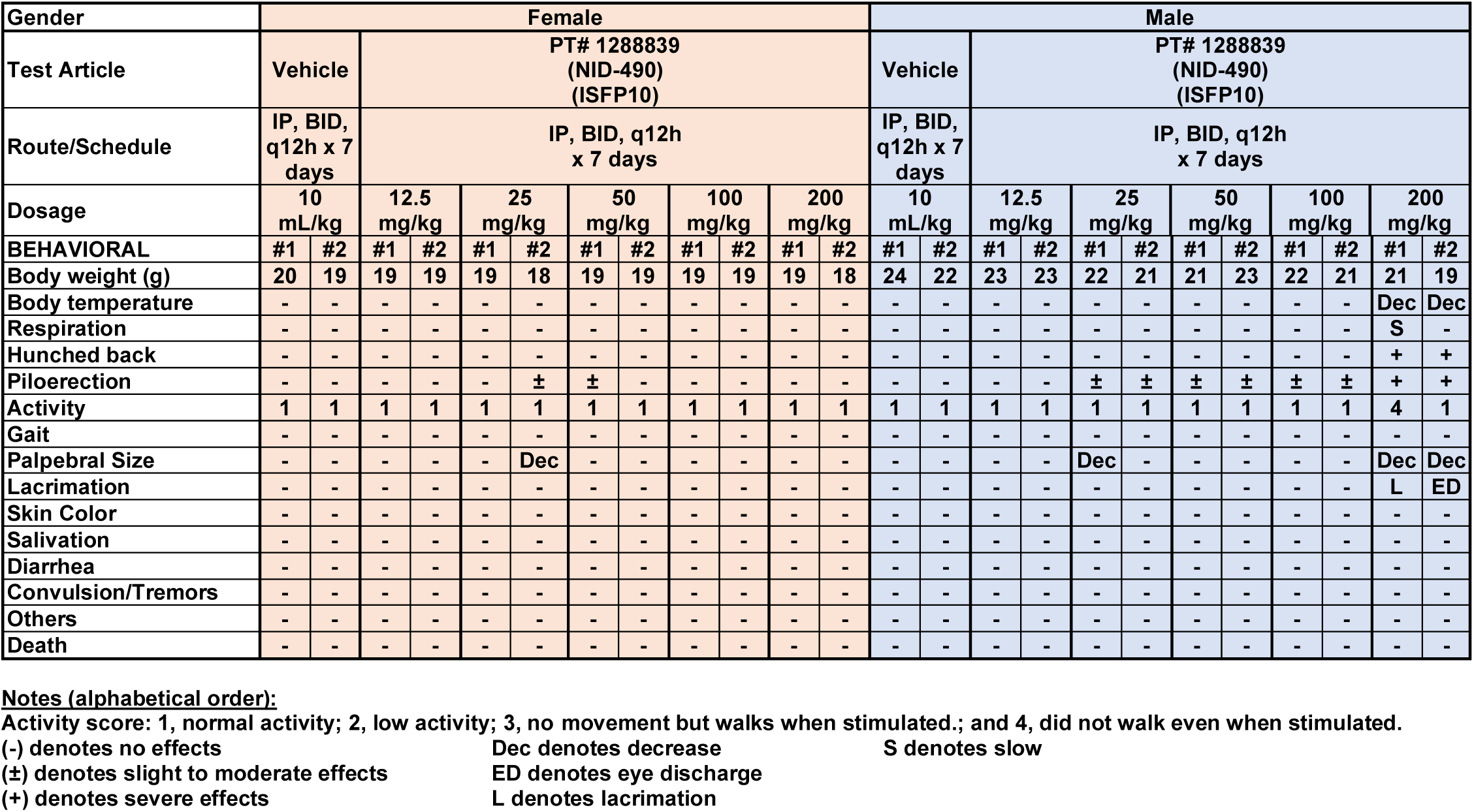
MTD results of ISFP10, cage side observation at 30 minutes after the 3^rd^ IP dosing.

| Gender | Female |  |  |  |  |  |  |  |  |  |  |  | Male |  |  |  |  |  |  |  |  |  |  |  |
| --- | --- | --- | --- | --- | --- | --- | --- | --- | --- | --- | --- | --- | --- | --- | --- | --- | --- | --- | --- | --- | --- | --- | --- | --- |
| Test Article | Vehicle |  | PT# 1288839<br>(NID-490)<br>(ISFP10) |  |  |  |  |  |  |  |  |  | Vehicle |  | PT# 1288839<br>(NID-490)<br>(ISFP10) |  |  |  |  |  |  |  |  |  |
| Route/Schedule | IP, BID,<br>q12h x 7<br>days |  | IP, BID, q12h<br>x 7 days |  |  |  |  |  |  |  |  |  | IP, BID,<br>q12h x 7<br>days |  | IP, BID, q12h<br>x 7 days |  |  |  |  |  |  |  |  |  |
| Dosage | 10<br>mL/kg |  | 12.5<br>mg/kg |  | 25<br>mg/kg |  | 50<br>mg/kg |  | 100<br>mg/kg |  | 200<br>mg/kg |  | 10<br>mL/kg |  | 12.5<br>mg/kg |  | 25<br>mg/kg |  | 50<br>mg/kg |  | 100<br>mg/kg |  | 200<br>mg/kg |  |
| BEHAVIORAL | #1 | #2 | #1 | #2 | #1 | #2 | #1 | #2 | #1 | #2 | #1 | #2 | #1 | #2 | #1 | #2 | #1 | #2 | #1 | #2 | #1 | #2 | #1 | #2 |
| Body weight (g) | 20 | 19 | 19 | 19 | 19 | 18 | 19 | 19 | 19 | 19 | 19 | 18 | 24 | 22 | 23 | 23 | 22 | 21 | 21 | 23 | 22 | 21 | 21 | 19 |
| Body temperature | - | - | - | - | - | - | - | - | - | - | - | - | - | - | - | - | - | - | - | - | - | - | Dec | Dec |
| Respiration | - | - | - | - | - | - | - | - | - | - | - | - | - | - | - | - | - | - | - | - | - | - | S | - |
| Hunched back | - | - | - | - | - | - | - | - | - | - | - | - | - | - | - | - | - | - | - | - | - | - | + | + |
| Piloerection | - | - | - | - | - | ± | ± | - | - | - | - | - | - | - | - | - | ± | ± | ± | ± | ± | ± | + | + |
| Activity | 1 | 1 | 1 | 1 | 1 | 1 | 1 | 1 | 1 | 1 | 1 | 1 | 1 | 1 | 1 | 1 | 1 | 1 | 1 | 1 | 1 | 1 | 4 | 1 |
| Gait | - | - | - | - | - | - | - | - | - | - | - | - | - | - | - | - | - | - | - | - | - | - | - | - |
| Palpebral Size | - | - | - | - | - | Dec | - | - | - | - | - | - | - | - | - | - | Dec | - | - | - | - | - | Dec | Dec |
| Lacrimation | - | - | - | - | - | - | - | - | - | - | - | - | - | - | - | - | - | - | - | - | - | - | L | ED |
| Skin Color | - | - | - | - | - | - | - | - | - | - | - | - | - | - | - | - | - | - | - | - | - | - | - | - |
| Salivation | - | - | - | - | - | - | - | - | - | - | - | - | - | - | - | - | - | - | - | - | - | - | - | - |
| Diarrhea | - | - | - | - | - | - | - | - | - | - | - | - | - | - | - | - | - | - | - | - | - | - | - | - |
| Convulsion/Tremors | - | - | - | - | - | - | - | - | - | - | - | - | - | - | - | - | - | - | - | - | - | - | - | - |
| Others | - | - | - | - | - | - | - | - | - | - | - | - | - | - | - | - | - | - | - | - | - | - | - | - |
| Death | - | - | - | - | - | - | - | - | - | - | - | - | - | - | - | - | - | - | - | - | - | - | - | - |

**Notes (alphabetical order):**

**Table 3-5.**
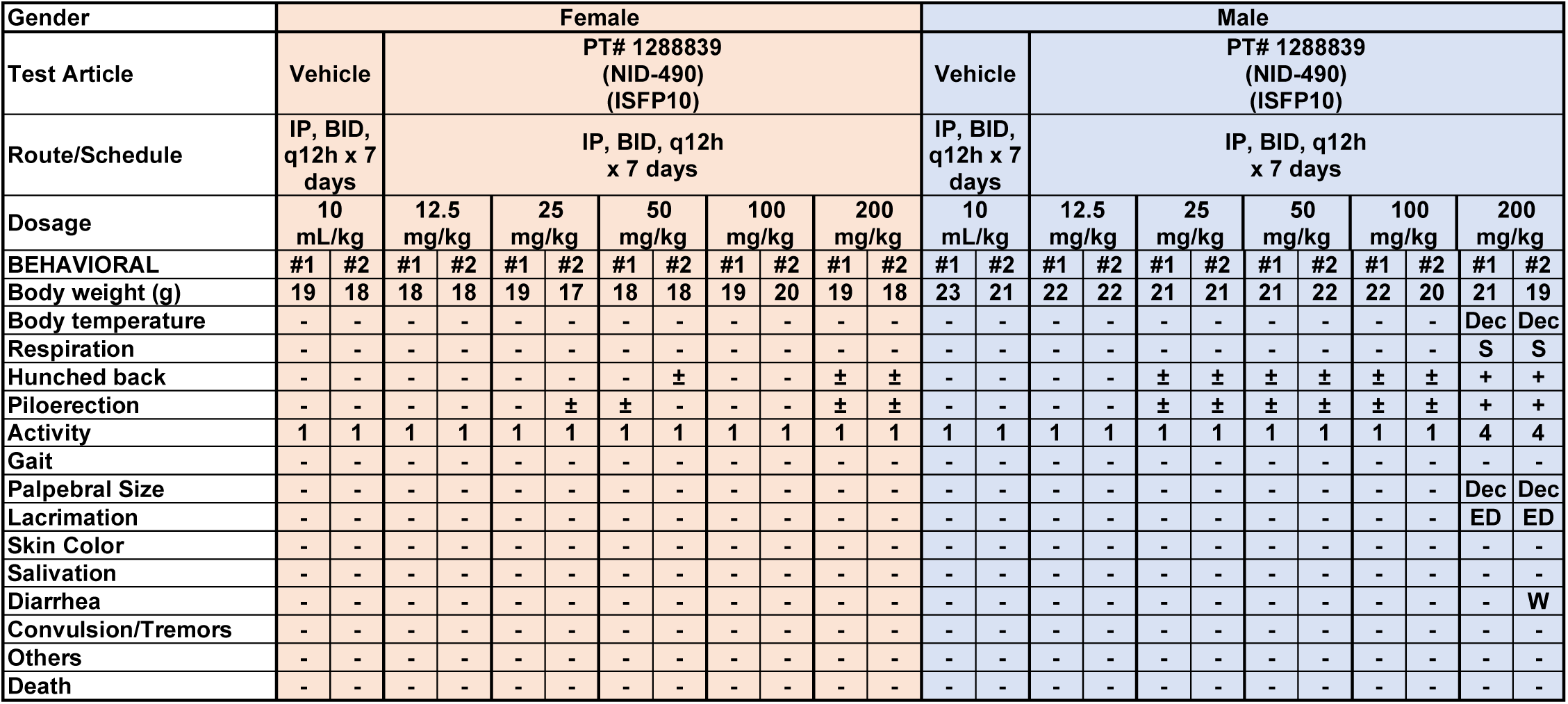
MTD results of ISFP10, cage side observation at 30 minutes after the 4^th^ IP dosing.

**Table 3-6.** MTD results of ISFP10, cage side observation at 30 minutes after the 5^th^ IP dosing.

| Gender | Female |  |  |  |  |  |  |  |  |  |  |  | Male |  |  |  |  |  |  |  |  |  |  |  |
| --- | --- | --- | --- | --- | --- | --- | --- | --- | --- | --- | --- | --- | --- | --- | --- | --- | --- | --- | --- | --- | --- | --- | --- | --- |
| Test Article | Vehicle |  | PT# 1288839<br>(NID-490)<br>(ISFP10) |  |  |  |  |  |  |  |  |  | Vehicle |  | PT# 1288839<br>(NID-490)<br>(ISFP10) |  |  |  |  |  |  |  |  |  |
| Route/Schedule | IP, BID,<br>q12h x 7<br>days |  | IP, BID, q12h<br>x 7 days |  |  |  |  |  |  |  |  |  | IP, BID,<br>q12h x 7<br>days |  | IP, BID, q12h<br>x 7 days |  |  |  |  |  |  |  |  |  |
| Dosage | 10<br>mL/kg |  | 12.5<br>mg/kg |  | 25<br>mg/kg |  | 50<br>mg/kg |  | 100<br>mg/kg |  | 200<br>mg/kg |  | 10<br>mL/kg |  | 12.5<br>mg/kg |  | 25<br>mg/kg |  | 50<br>mg/kg |  | 100<br>mg/kg |  | 200<br>mg/kg |  |
| BEHAVIORAL | #1 | #2 | #1 | #2 | #1 | #2 | #1 | #2 | #1 | #2 | #1 | #2 | #1 | #2 | #1 | #2 | #1 | #2 | #1 | #2 | #1 | #2 | #1 | #2 |
| Body weight (g) | 21 | 19 | 19 | 19 | 20 | 17 | 18 | 19 | 20 | 20 | 18 | 18 | 24 | 22 | 23 | 23 | 22 | 22 | 22 | 23 | 22 | 21 |  |  |
| Body temperature | - | - | - | - | - | - | - | - | - | - | - | - | - | - | - | - | - | - | - | - | - | - |  |  |
| Respiration | - | - | - | - | - | - | - | - | - | - | - | - | - | - | - | - | - | - | - | - | - | - |  |  |
| Hunched back | - | - | - | - | - | - | ± | - | - | - | ± | ± | - | - | - | - | ± | ± | ± | ± | ± | ± |  |  |
| Piloerection | - | - | - | - | - | ± | ± | - | - | - | ± | ± | - | - | - | - | ± | ± | ± | ± | ± | ± |  |  |
| Activity | 1 | 1 | 1 | 1 | 1 | 1 | 1 | 1 | 1 | 1 | 1 | 1 | 1 | 1 | 1 | 1 | 1 | 1 | 1 | 1 | 1 | 1 |  |  |
| Gait | - | - | - | - | - | - | - | - | - | - | - | - | - | - | - | - | - | - | - | - | - | - |  |  |
| Palpebral Size | - | - | - | - | - | - | - | - | - | - | - | - | - | - | - | - | - | - | - | - | - | - |  |  |
| Lacrimation | - | - | - | - | - | - | - | - | - | - | - | - | - | - | - | - | - | - | - | - | - | - |  |  |
| Skin Color | - | - | - | - | - | - | - | - | - | - | - | - | - | - | - | - | - | - | - | - | - | - |  |  |
| Salivation | - | - | - | - | - | - | - | - | - | - | - | - | - | - | - | - | - | - | - | - | - | - |  |  |
| Diarrhea | - | - | - | - | - | - | - | - | - | - | - | - | - | - | - | - | - | - | - | - | - | W |  |  |
| Convulsion/Tremors | - | - | - | - | - | - | - | - | - | - | - | - | - | - | - | - | - | - | - | - | - | - |  |  |
| Others | - | - | - | - | - | - | - | - | - | - | - | - | - | - | - | - | - | - | - | - | - | - |  |  |
| Death | - | - | - | - | - | - | - | - | - | - | - | - | - | - | - | - | - | - | - | - | - | - | Dead | Dead |

**Notes (alphabetical order):**
Activity score: 1, normal activity; 2, low activity; 3, no movement but walks when stimulated.; and 4, did not walk even when stimulated.
(-) denotes no effects
Dec denotes decrease
W denotes wet tail
(±) denotes slight to moderate effects
ED denotes eye discharge
(+) denotes severe effects
S denotes slow

**Table 3-7.**
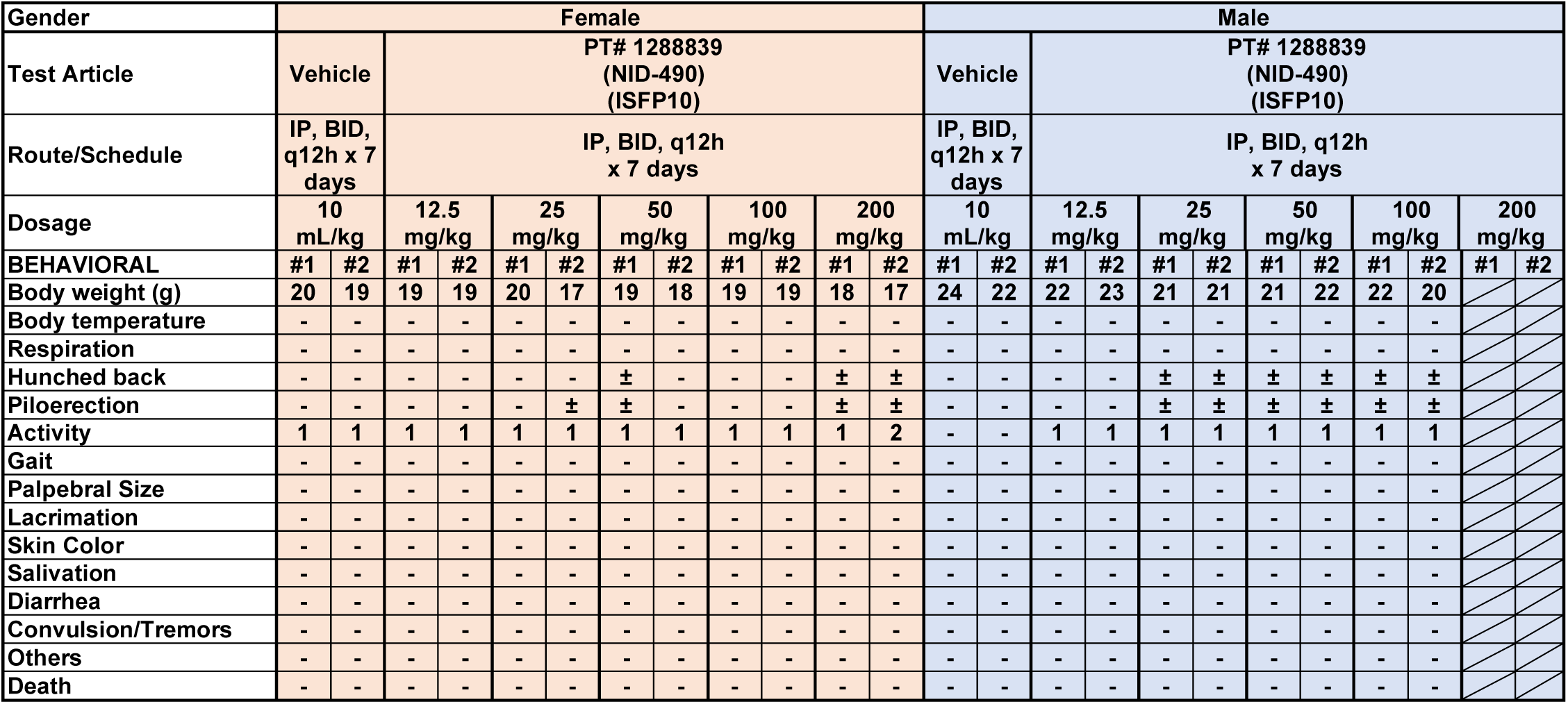
MTD results of ISFP10, cage side observation at 30 minutes after the 6^th^ IP dosing.

**Table 3-8.** MTD results of ISFP10, cage side observation at 30 minutes after the 7^th^ IP dosing.

| Gender | Female |  |  |  |  |  |  |  |  |  |  |  | Male |  |  |  |  |  |  |  |  |  |  |  |
| --- | --- | --- | --- | --- | --- | --- | --- | --- | --- | --- | --- | --- | --- | --- | --- | --- | --- | --- | --- | --- | --- | --- | --- | --- |
| Test Article | Vehicle |  | PT# 1288839<br>(NID-490)<br>(ISFP10) |  |  |  |  |  |  |  |  |  | Vehicle |  | PT# 1288839<br>(NID-490)<br>(ISFP10) |  |  |  |  |  |  |  |  |  |
| Route/Schedule | IP, BID,<br>q12h x 7<br>days |  | IP, BID, q12h<br>x 7 days |  |  |  |  |  |  |  |  |  | IP, BID,<br>q12h x 7<br>days |  | IP, BID, q12h<br>x 7 days |  |  |  |  |  |  |  |  |  |
| Dosage | 10<br>mL/kg |  | 12.5<br>mg/kg |  | 25<br>mg/kg |  | 50<br>mg/kg |  | 100<br>mg/kg |  | 200<br>mg/kg |  | 10<br>mL/kg |  | 12.5<br>mg/kg |  | 25<br>mg/kg |  | 50<br>mg/kg |  | 100<br>mg/kg |  | 200<br>mg/kg |  |
| BEHAVIORAL | #1 | #2 | #1 | #2 | #1 | #2 | #1 | #2 | #1 | #2 | #1 | #2 | #1 | #2 | #1 | #2 | #1 | #2 | #1 | #2 | #1 | #2 | #1 | #2 |
| Body weight (g) | 22 | 20 | 19 | 19 | 21 | 18 | 20 | 20 | 20 | 20 | 18 | 17 | 24 | 23 | 24 | 24 | 23 | 22 | 23 | 23 | 21 | 20 |  |  |
| Body temperature | - | - | - | - | - | - | - | - | - | - | - | Dec | - | - | - | - | - | - | - | - | - | - |  |  |
| Respiration | - | - | - | - | - | - | - | - | - | - | - | S | - | - | - | - | - | - | - | - | - | - |  |  |
| Hunched back | - | - | - | - | - | - | - | - | - | - | ± | + | - | - | - | - | ± | ± | ± | ± | ± | ± |  |  |
| Piloerection | - | - | - | - | - | ± | ± | - | - | - | ± | + | - | - | - | - | ± | ± | ± | ± | ± | ± |  |  |
| Activity | 1 | 1 | 1 | 1 | 1 | 1 | 1 | 1 | 1 | 1 | 1 | 4 | - | - | 1 | 1 | 1 | 1 | 1 | 1 | 1 | 1 |  |  |
| Gait | - | - | - | - | - | - | - | - | - | - | - | - | - | - | - | - | - | - | - | - | - | - |  |  |
| Palpebral Size | - | - | - | - | - | - | - | - | - | - | - | - | - | - | - | - | - | - | - | - | - | - |  |  |
| Lacrimation | - | - | - | - | - | - | - | - | - | - | - | - | - | - | - | - | - | - | - | - | - | - |  |  |
| Skin Color | - | - | - | - | - | - | - | - | - | - | - | - | - | - | - | - | - | - | - | - | - | - |  |  |
| Salivation | - | - | - | - | - | - | - | - | - | - | - | - | - | - | - | - | - | - | - | - | - | - |  |  |
| Diarrhea | - | - | - | - | - | - | - | - | - | - | - | W | - | - | - | - | - | - | - | - | - | - |  |  |
| Convulsion/Tremors | - | - | - | - | - | - | - | - | - | - | - | - | - | - | - | - | - | - | - | - | - | - |  |  |
| Others | - | - | - | - | - | - | - | - | - | - | - | - | - | - | - | - | - | - | - | - | - | - |  |  |
| Death | - | - | - | - | - | - | - | - | - | - | - | - | - | - | - | - | - | - | - | - | - | - |  |  |

**Notes (alphabetical order):**

**Table 3-9.**
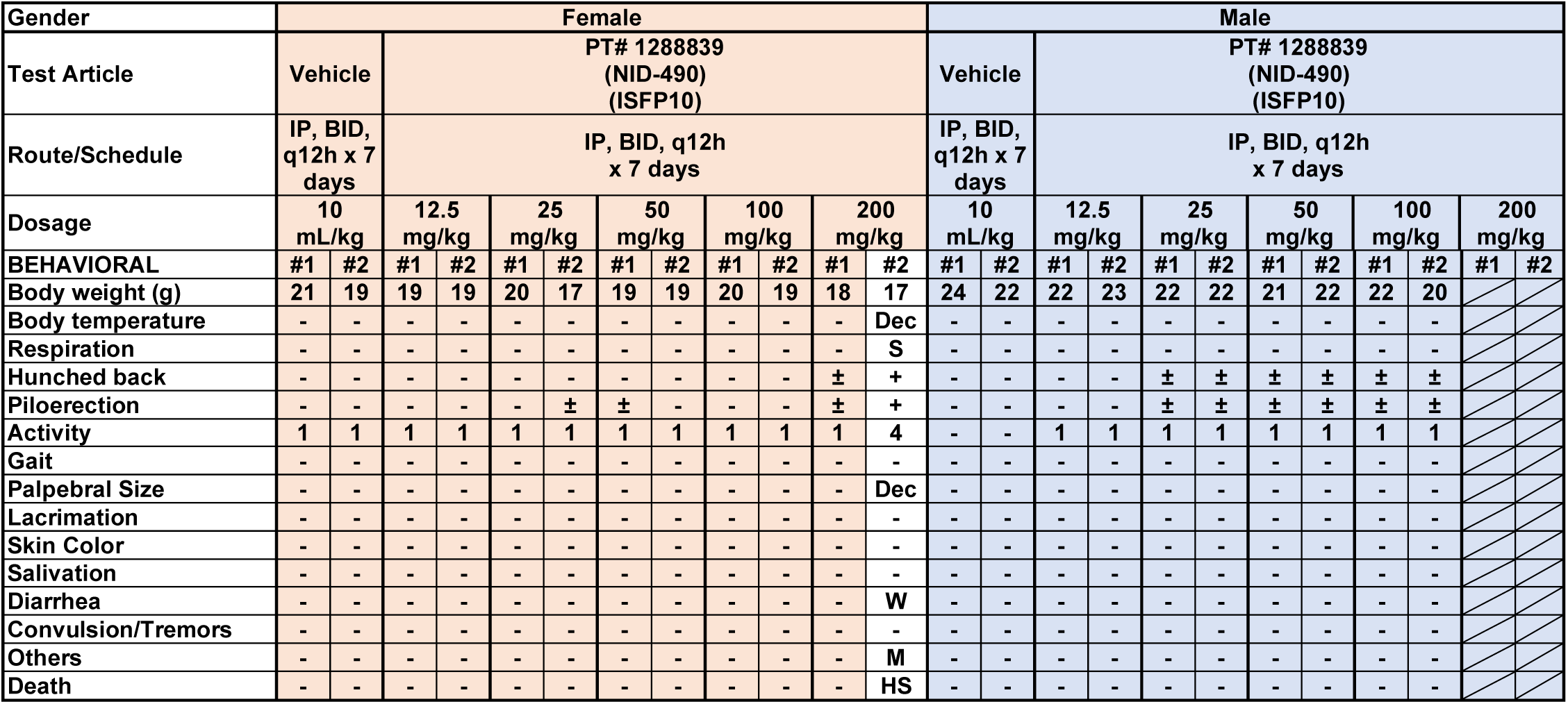
MTD results of ISFP10, cage side observation at 30 minutes after the 8^th^ IP dosing.

**Table 3-10.** MTD results of ISFP10, cage side observation at 30 minutes after the 9^th^ IP dosing.

| Gender | Female |  |  |  |  |  |  |  |  |  |  |  | Male |  |  |  |  |  |  |  |  |  |  |  |
| --- | --- | --- | --- | --- | --- | --- | --- | --- | --- | --- | --- | --- | --- | --- | --- | --- | --- | --- | --- | --- | --- | --- | --- | --- |
| Test Article | Vehicle |  | PT# 1288839<br>(NID-490)<br>(ISFP10) |  |  |  |  |  |  |  |  |  | Vehicle |  | PT# 1288839<br>(NID-490)<br>(ISFP10) |  |  |  |  |  |  |  |  |  |
| Route/Schedule | IP, BID,<br>q12h x 7<br>days |  | IP, BID, q12h<br>x 7 days |  |  |  |  |  |  |  |  |  | IP, BID,<br>q12h x 7<br>days |  | IP, BID, q12h<br>x 7 days |  |  |  |  |  |  |  |  |  |
| Dosage | 10<br>mL/kg |  | 12.5<br>mg/kg |  | 25<br>mg/kg |  | 50<br>mg/kg |  | 100<br>mg/kg |  | 200<br>mg/kg |  | 10<br>mL/kg |  | 12.5<br>mg/kg |  | 25<br>mg/kg |  | 50<br>mg/kg |  | 100<br>mg/kg |  | 200<br>mg/kg |  |
| BEHAVIORAL | #1 | #2 | #1 | #2 | #1 | #2 | #1 | #2 | #1 | #2 | #1 | #2 | #1 | #2 | #1 | #2 | #1 | #2 | #1 | #2 | #1 | #2 | #1 | #2 |
| Body weight (g) | 22 | 20 | 20 | 19 | 21 | 19 | 20 | 20 | 21 | 19 | 18 |  | 25 | 23 | 23 | 24 | 23 | 23 | 22 | 23 | 22 | 21 |  |  |
| Body temperature | - | - | - | - | - | - | - | - | - | - | - |  | - | - | - | - | - | - | - | - | - | - |  |  |
| Respiration | - | - | - | - | - | - | - | - | - | - | - |  | - | - | - | - | - | - | - | - | - | - |  |  |
| Hunched back | - | - | - | - | - | - | - | - | - | - | ± |  | - | - | - | - | ± | ± | ± | ± | ± | ± |  |  |
| Piloerection | - | - | - | - | - | ± | ± | - | - | - | ± |  | - | - | - | - | ± | ± | ± | ± | ± | ± |  |  |
| Activity | 1 | 1 | 1 | 1 | 1 | 1 | 1 | 1 | 1 | 1 | 1 |  | - | - | 1 | 1 | 1 | 1 | 1 | 1 | 1 | 1 |  |  |
| Gait | - | - | - | - | - | - | - | - | - | - | - |  | - | - | - | - | - | - | - | - | - | - |  |  |
| Palpebral Size | - | - | - | - | - | - | - | - | - | - | - |  | - | - | - | - | - | - | - | - | - | - |  |  |
| Lacrimation | - | - | - | - | - | - | - | - | - | - | - |  | - | - | - | - | - | - | - | - | - | - |  |  |
| Skin Color | - | - | - | - | - | - | - | - | - | - | - |  | - | - | - | - | - | - | - | - | - | - |  |  |
| Salivation | - | - | - | - | - | - | - | - | - | - | - |  | - | - | - | - | - | - | - | - | - | - |  |  |
| Diarrhea | - | - | - | - | - | - | - | - | - | - | - |  | - | - | - | - | - | - | - | - | - | - |  |  |
| Convulsion/Tremors | - | - | - | - | - | - | - | - | - | - | - |  | - | - | - | - | - | - | - | - | - | - |  |  |
| Others | - | - | - | - | - | - | - | - | - | - | - |  | - | - | - | - | - | - | - | - | - | - |  |  |
| Death | - | - | - | - | - | - | - | - | - | - | - |  | - | - | - | - | - | - | - | - | - | - |  |  |

**Notes (alphabetical order):**

**Table 3-11.** MTD results of ISFP10, cage side observation at 30 minutes after the 10^th^ IP dosing.

| Gender | Female |  |  |  |  |  |  |  |  |  |  |  | Male |  |  |  |  |  |  |  |  |  |  |  |
| --- | --- | --- | --- | --- | --- | --- | --- | --- | --- | --- | --- | --- | --- | --- | --- | --- | --- | --- | --- | --- | --- | --- | --- | --- |
| Test Article | Vehicle |  | PT# 1288839<br>(NID-490)<br>(ISFP10) |  |  |  |  |  |  |  |  |  | Vehicle |  | PT# 1288839<br>(NID-490)<br>(ISFP10) |  |  |  |  |  |  |  |  |  |
| Route/Schedule | IP, BID,<br>q12h x 7<br>days |  | IP, BID, q12h<br>x 7 days |  |  |  |  |  |  |  |  |  | IP, BID,<br>q12h x 7<br>days |  | IP, BID, q12h<br>x 7 days |  |  |  |  |  |  |  |  |  |
| Dosage | 10<br>mL/kg |  | 12.5<br>mg/kg |  | 25<br>mg/kg |  | 50<br>mg/kg |  | 100<br>mg/kg |  | 200<br>mg/kg |  | 10<br>mL/kg |  | 12.5<br>mg/kg |  | 25<br>mg/kg |  | 50<br>mg/kg |  | 100<br>mg/kg |  | 200<br>mg/kg |  |
| BEHAVIORAL | #1 | #2 | #1 | #2 | #1 | #2 | #1 | #2 | #1 | #2 | #1 | #2 | #1 | #2 | #1 | #2 | #1 | #2 | #1 | #2 | #1 | #2 | #1 | #2 |
| Body weight (g) | 21 | 19 | 20 | 19 | 20 | 18 | 19 | 19 | 20 | 19 | 18 |  | 23 | 22 | 21 | 23 | 22 | 21 | 21 | 22 | 21 | 20 |  |  |
| Body temperature | - | - | - | - | - | - | - | - | - | - | - |  | - | - | - | - | - | - | - | - | - | - |  |  |
| Respiration | - | - | - | - | - | - | - | - | - | - | - |  | - | - | - | - | - | - | - | - | - | - |  |  |
| Hunched back | - | - | - | - | - | - | ± | - | - | - | ± |  | - | - | - | ± | ± | ± | ± | ± | ± | ± |  |  |
| Piloerection | - | - | - | - | - | ± | ± | - | - | - | ± |  | - | - | - | ± | ± | ± | ± | ± | ± | ± |  |  |
| Activity | 1 | 1 | 1 | 1 | 1 | 1 | 1 | 1 | 1 | 1 | 1 |  | - | - | 1 | 1 | 1 | 1 | 1 | 1 | 1 | 1 |  |  |
| Gait | - | - | - | - | - | - | - | - | - | - | - |  | - | - | - | - | - | - | - | - | - | - |  |  |
| Palpebral Size | - | - | - | - | - | - | - | - | - | - | - |  | - | - | - | - | - | - | - | - | - | - |  |  |
| Lacrimation | - | - | - | - | - | - | - | - | - | - | - |  | - | - | - | - | - | - | - | - | - | - |  |  |
| Skin Color | - | - | - | - | - | - | - | - | - | - | - |  | - | - | - | - | - | - | - | - | - | - |  |  |
| Salivation | - | - | - | - | - | - | - | - | - | - | - |  | - | - | - | - | - | - | - | - | - | - |  |  |
| Diarrhea | - | - | - | - | - | - | - | - | - | - | - |  | - | - | - | - | - | - | - | - | - | - |  |  |
| Convulsion/Tremors | - | - | - | - | - | - | - | - | - | - | - |  | - | - | - | - | - | - | - | - | - | - |  |  |
| Others | - | - | - | - | - | - | - | - | - | - | - |  | - | - | - | - | - | - | - | - | - | - |  |  |
| Death | - | - | - | - | - | - | - | - | - | - | - |  | - | - | - | - | - | - | - | - | - | - |  |  |

**Table 3-12.** MTD results of ISFP10, cage side observation at 30 minutes after the 11^th^ IP dosing.

| Gender | Female |  |  |  |  |  |  |  |  |  |  |  | Male |  |  |  |  |  |  |  |  |  |  |  |
| --- | --- | --- | --- | --- | --- | --- | --- | --- | --- | --- | --- | --- | --- | --- | --- | --- | --- | --- | --- | --- | --- | --- | --- | --- |
| Test Article | Vehicle |  | PT# 1288839<br>(NID-490)<br>(ISFP10) |  |  |  |  |  |  |  |  |  | Vehicle |  | PT# 1288839<br>(NID-490)<br>(ISFP10) |  |  |  |  |  |  |  |  |  |
| Route/Schedule | IP, BID,<br>q12h x 7<br>days |  | IP, BID, q12h<br>x 7 days |  |  |  |  |  |  |  |  |  | IP, BID,<br>q12h x 7<br>days |  | IP, BID, q12h<br>x 7 days |  |  |  |  |  |  |  |  |  |
| Dosage | 10<br>mL/kg |  | 12.5<br>mg/kg |  | 25<br>mg/kg |  | 50<br>mg/kg |  | 100<br>mg/kg |  | 200<br>mg/kg |  | 10<br>mL/kg |  | 12.5<br>mg/kg |  | 25<br>mg/kg |  | 50<br>mg/kg |  | 100<br>mg/kg |  | 200<br>mg/kg |  |
| BEHAVIORAL | #1 | #2 | #1 | #2 | #1 | #2 | #1 | #2 | #1 | #2 | #1 | #2 | #1 | #2 | #1 | #2 | #1 | #2 | #1 | #2 | #1 | #2 | #1 | #2 |
| Body weight (g) | 21 | 20 | 20 | 19 | 20 | 18 | 20 | 19 | 21 | 19 | 18 |  | 24 | 23 | 23 | 24 | 24 | 23 | 22 | 22 | 22 | 21 |  |  |
| Body temperature | - | - | - | - | - | - | - | - | - | - | - |  | - | - | - | - | - | - | - | - | - | - |  |  |
| Respiration | - | - | - | - | - | - | - | - | - | - | - |  | - | - | - | - | - | - | - | - | - | - |  |  |
| Hunched back | - | - | - | ± | - | - | - | - | - | ± | ± |  | - | - | - | ± | ± | ± | ± | ± | ± | ± |  |  |
| Piloerection | - | - | - | ± | - | ± | ± | - | - | ± | ± |  | - | - | - | ± | ± | ± | ± | ± | ± | ± |  |  |
| Activity | 1 | 1 | 1 | 1 | 1 | 1 | 1 | 1 | 1 | 1 | 1 |  | - | - | 1 | 1 | 1 | 1 | 1 | 1 | 1 | 1 |  |  |
| Gait | - | - | - | - | - | - | - | - | - | - | - |  | - | - | - | - | - | - | - | - | - | - |  |  |
| Palpebral Size | - | - | - | - | - | - | - | - | - | - | - |  | - | - | - | - | - | - | - | - | - | - |  |  |
| Lacrimation | - | - | - | - | - | - | - | - | - | - | - |  | - | - | - | - | - | - | - | - | - | - |  |  |
| Skin Color | - | - | - | - | - | - | - | - | - | - | - |  | - | - | - | - | - | - | - | - | - | - |  |  |
| Salivation | - | - | - | - | - | - | - | - | - | - | - |  | - | - | - | - | - | - | - | - | - | - |  |  |
| Diarrhea | - | - | - | - | - | - | - | - | - | - | - |  | - | - | - | - | - | - | - | - | - | - |  |  |
| Convulsion/Tremors | - | - | - | - | - | - | - | - | - | - | - |  | - | - | - | - | - | - | - | - | - | - |  |  |
| Others | - | - | - | - | - | - | - | - | - | - | - |  | - | - | - | - | - | - | - | - | - | - |  |  |
| Death | - | - | - | - | - | - | - | - | - | - | - |  | - | - | - | - | - | - | - | - | - | - |  |  |

**Notes (alphabetical order):**

**Table 3-13.**
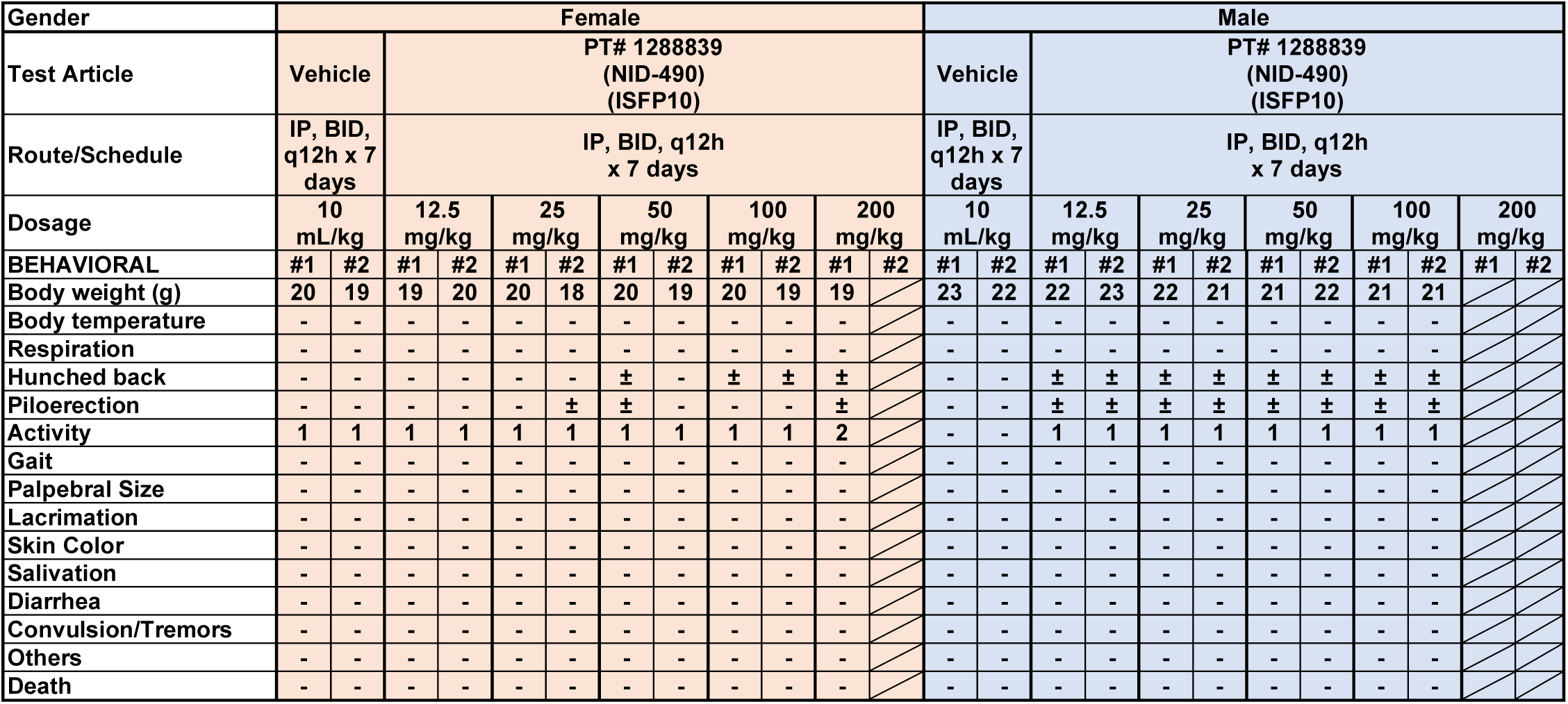
MTD results of ISFP10, cage side observation at 30 minutes after the 12^th^ IP dosing.

**Table 3-14.** MTD results of ISFP10, cage side observation at 30 minutes after the 13^th^ IP dosing.

| Gender | Female |  |  |  |  |  |  |  |  |  |  |  | Male |  |  |  |  |  |  |  |  |  |  |  |
| --- | --- | --- | --- | --- | --- | --- | --- | --- | --- | --- | --- | --- | --- | --- | --- | --- | --- | --- | --- | --- | --- | --- | --- | --- |
| Test Article | Vehicle |  | PT# 1288839<br>(NID-490)<br>(ISFP10) |  |  |  |  |  |  |  |  |  | Vehicle |  | PT# 1288839<br>(NID-490)<br>(ISFP10) |  |  |  |  |  |  |  |  |  |
| Route/Schedule | IP, BID,<br>q12h x 7<br>days |  | IP, BID, q12h<br>x 7 days |  |  |  |  |  |  |  |  |  | IP, BID,<br>q12h x 7<br>days |  | IP, BID, q12h<br>x 7 days |  |  |  |  |  |  |  |  |  |
| Dosage | 10<br>mL/kg |  | 12.5<br>mg/kg |  | 25<br>mg/kg |  | 50<br>mg/kg |  | 100<br>mg/kg |  | 200<br>mg/kg |  | 10<br>mL/kg |  | 12.5<br>mg/kg |  | 25<br>mg/kg |  | 50<br>mg/kg |  | 100<br>mg/kg |  | 200<br>mg/kg |  |
| BEHAVIORAL | #1 | #2 | #1 | #2 | #1 | #2 | #1 | #2 | #1 | #2 | #1 | #2 | #1 | #2 | #1 | #2 | #1 | #2 | #1 | #2 | #1 | #2 | #1 | #2 |
| Body weight (g) | 22 | 20 | 20 | 19 | 21 | 19 | 21 | 18 | 20 | 20 | 19 |  | 24 | 24 | 23 | 24 | 22 | 21 | 23 | 23 | 23 | 20 |  |  |
| Body temperature | - | - | - | - | - | - | - | - | - | - | - |  | - | - | - | - | - | - | - | - | - | - |  |  |
| Respiration | - | - | - | - | - | - | - | - | - | - | - |  | - | - | - | - | - | - | - | - | - | - |  |  |
| Hunched back | - | - | - | - | - | - | ± | ± | ± | ± | ± |  | - | - | ± | ± | ± | ± | ± | ± | ± | ± |  |  |
| Piloerection | - | - | - | - | - | ± | ± | ± | ± | ± | ± |  | - | - | ± | ± | ± | ± | ± | ± | ± | ± |  |  |
| Activity | 1 | 1 | 1 | 1 | 1 | 1 | 1 | 1 | 1 | 1 | 1 |  | - | - | 1 | 1 | 1 | 1 | 1 | 1 | 1 | 1 |  |  |
| Gait | - | - | - | - | - | - | - | - | - | - | - |  | - | - | - | - | - | - | - | - | - | - |  |  |
| Palpebral Size | - | - | - | - | - | - | - | - | - | - | - |  | - | - | - | - | - | - | - | - | - | - |  |  |
| Lacrimation | - | - | - | - | - | - | - | - | - | - | - |  | - | - | - | - | - | - | - | - | - | - |  |  |
| Skin Color | - | - | - | - | - | - | - | - | - | - | - |  | - | - | - | - | - | - | - | - | - | - |  |  |
| Salivation | - | - | - | - | - | - | - | - | - | - | - |  | - | - | - | - | - | - | - | - | - | - |  |  |
| Diarrhea | - | - | - | - | - | - | - | - | - | - | - |  | - | - | - | - | - | - | - | - | - | - |  |  |
| Convulsion/Tremors | - | - | - | - | - | - | - | - | - | - | - |  | - | - | - | - | - | - | - | - | - | - |  |  |
| Others | - | - | - | - | - | - | - | - | - | - | - |  | - | - | - | - | - | - | - | - | - | - |  |  |
| Death | - | - | - | - | - | - | - | - | - | - | - |  | - | - | - | - | - | - | - | - | - | - |  |  |

**Notes (alphabetical order):**

**Table 3-15.**
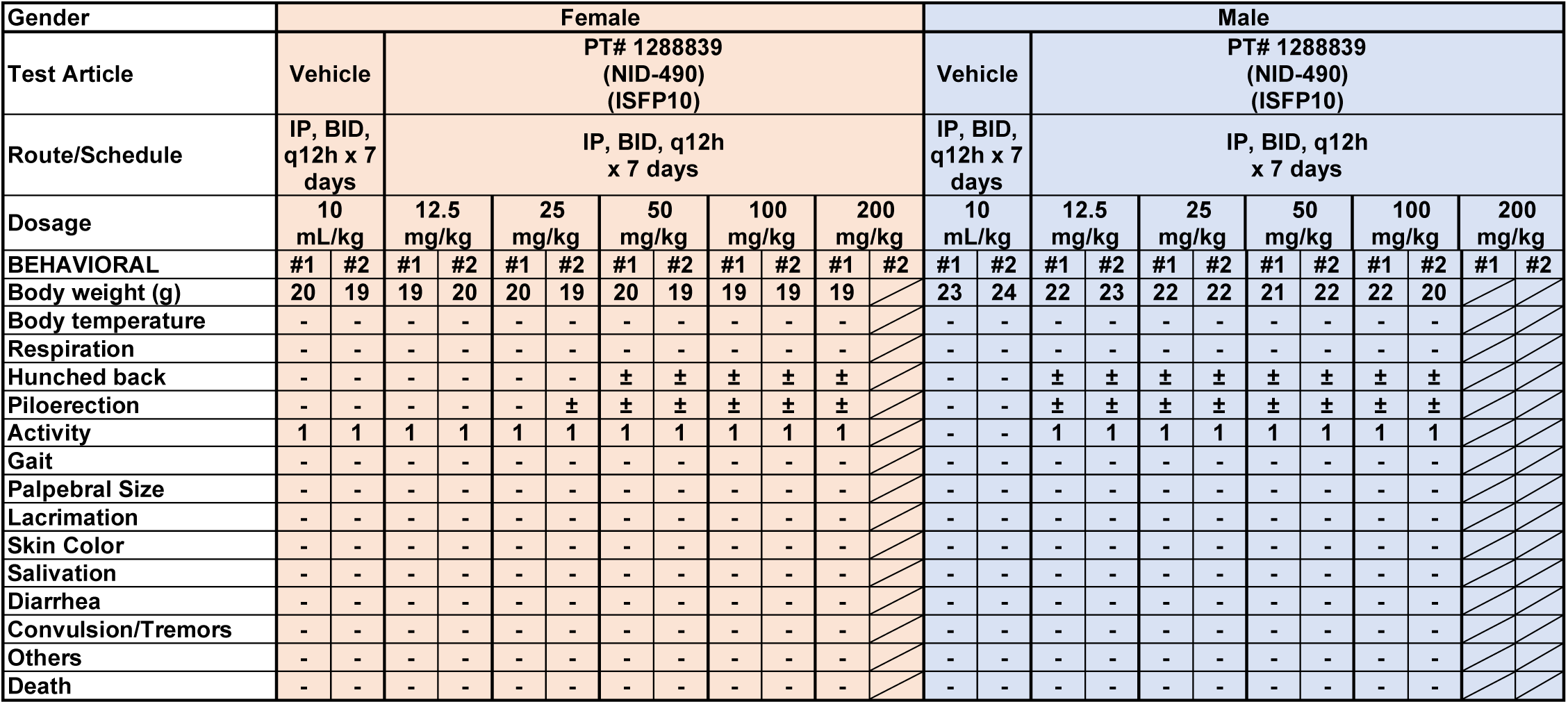
MTD results of ISFP10, cage side observation at 30 minutes after the 14^th^ IP dosing.

| Gender | Female |  |  |  |  |  |  |  |  |  |  |  | Male |  |  |  |  |  |  |  |  |  |  |  |
| --- | --- | --- | --- | --- | --- | --- | --- | --- | --- | --- | --- | --- | --- | --- | --- | --- | --- | --- | --- | --- | --- | --- | --- | --- |
| Test Article | Vehicle |  | PT# 1288839<br>(NID-490)<br>(ISFP10) |  |  |  |  |  |  |  |  |  | Vehicle |  | PT# 1288839<br>(NID-490)<br>(ISFP10) |  |  |  |  |  |  |  |  |  |
| Route/Schedule | IP, BID,<br>q12h x 7<br>days |  | IP, BID, q12h<br>x 7 days |  |  |  |  |  |  |  |  |  | IP, BID,<br>q12h x 7<br>days |  | IP, BID, q12h<br>x 7 days |  |  |  |  |  |  |  |  |  |
| Dosage | 10<br>mL/kg |  | 12.5<br>mg/kg |  | 25<br>mg/kg |  | 50<br>mg/kg |  | 100<br>mg/kg |  | 200<br>mg/kg |  | 10<br>mL/kg |  | 12.5<br>mg/kg |  | 25<br>mg/kg |  | 50<br>mg/kg |  | 100<br>mg/kg |  | 200<br>mg/kg |  |
| BEHAVIORAL | #1 | #2 | #1 | #2 | #1 | #2 | #1 | #2 | #1 | #2 | #1 | #2 | #1 | #2 | #1 | #2 | #1 | #2 | #1 | #2 | #1 | #2 | #1 | #2 |
| Body weight (g) | 20 | 19 | 19 | 20 | 20 | 19 | 20 | 19 | 19 | 19 | 19 |  | 23 | 24 | 22 | 23 | 22 | 22 | 21 | 22 | 22 | 20 |  |  |
| Body temperature | - | - | - | - | - | - | - | - | - | - | - |  | - | - | - | - | - | - | - | - | - | - |  |  |
| Respiration | - | - | - | - | - | - | - | - | - | - | - |  | - | - | - | - | - | - | - | - | - | - |  |  |
| Hunched back | - | - | - | - | - | - | ± | ± | ± | ± | ± |  | - | - | ± | ± | ± | ± | ± | ± | ± | ± |  |  |
| Piloerection | - | - | - | - | - | ± | ± | ± | ± | ± | ± |  | - | - | ± | ± | ± | ± | ± | ± | ± | ± |  |  |
| Activity | 1 | 1 | 1 | 1 | 1 | 1 | 1 | 1 | 1 | 1 | 1 |  | - | - | 1 | 1 | 1 | 1 | 1 | 1 | 1 | 1 |  |  |
| Gait | - | - | - | - | - | - | - | - | - | - | - |  | - | - | - | - | - | - | - | - | - | - |  |  |
| Palpebral Size | - | - | - | - | - | - | - | - | - | - | - |  | - | - | - | - | - | - | - | - | - | - |  |  |
| Lacrimation | - | - | - | - | - | - | - | - | - | - | - |  | - | - | - | - | - | - | - | - | - | - |  |  |
| Skin Color | - | - | - | - | - | - | - | - | - | - | - |  | - | - | - | - | - | - | - | - | - | - |  |  |
| Salivation | - | - | - | - | - | - | - | - | - | - | - |  | - | - | - | - | - | - | - | - | - | - |  |  |
| Diarrhea | - | - | - | - | - | - | - | - | - | - | - |  | - | - | - | - | - | - | - | - | - | - |  |  |
| Convulsion/Tremors | - | - | - | - | - | - | - | - | - | - | - |  | - | - | - | - | - | - | - | - | - | - |  |  |
| Others | - | - | - | - | - | - | - | - | - | - | - |  | - | - | - | - | - | - | - | - | - | - |  |  |
| Death | - | - | - | - | - | - | - | - | - | - | - |  | - | - | - | - | - | - | - | - | - | - |  |  |

**Table 3-16.** MTD results of ISFP10, cage side observation at the 1^st^ day after the last dosing.

| Gender | Female |  |  |  |  |  |  |  |  |  |  |  | Male |  |  |  |  |  |  |  |  |  |  |  |
| --- | --- | --- | --- | --- | --- | --- | --- | --- | --- | --- | --- | --- | --- | --- | --- | --- | --- | --- | --- | --- | --- | --- | --- | --- |
| Test Article | Vehicle |  | PT# 1288839<br>(NID-490)<br>(ISFP10) |  |  |  |  |  |  |  |  |  | Vehicle |  | PT# 1288839<br>(NID-490)<br>(ISFP10) |  |  |  |  |  |  |  |  |  |
| Route/Schedule | IP, BID,<br>q12h x 7<br>days |  | IP, BID, q12h<br>x 7 days |  |  |  |  |  |  |  |  |  | IP, BID,<br>q12h x 7<br>days |  | IP, BID, q12h<br>x 7 days |  |  |  |  |  |  |  |  |  |
| Dosage | 10<br>mL/kg |  | 12.5<br>mg/kg |  | 25<br>mg/kg |  | 50<br>mg/kg |  | 100<br>mg/kg |  | 200<br>mg/kg |  | 10<br>mL/kg |  | 12.5<br>mg/kg |  | 25<br>mg/kg |  | 50<br>mg/kg |  | 100<br>mg/kg |  | 200<br>mg/kg |  |
| BEHAVIORAL | #1 | #2 | #1 | #2 | #1 | #2 | #1 | #2 | #1 | #2 | #1 | #2 | #1 | #2 | #1 | #2 | #1 | #2 | #1 | #2 | #1 | #2 | #1 | #2 |
| Body weight (g) | 22 | 20 | 20 | 21 | 21 | 20 | 20 | 19 | 19 | 20 | 19 |  | 24 | 25 | 24 | 24 | 23 | 23 | 23 | 23 | 22 | 21 |  |  |
| Body temperature | - | - | - | - | - | - | - | - | - | - | - |  | - | - | - | - | - | - | - | - | - | - |  |  |
| Respiration | - | - | - | - | - | - | - | - | - | - | - |  | - | - | - | - | - | - | - | - | - | - |  |  |
| Hunched back | - | - | - | - | - | - | ± | ± | ± | ± | ± |  | - | - | ± | ± | ± | ± | ± | ± | ± | ± |  |  |
| Piloerection | - | - | - | - | - | ± | ± | ± | ± | ± | ± |  | - | - | ± | ± | ± | ± | ± | ± | ± | ± |  |  |
| Activity | 1 | 1 | 1 | 1 | 1 | 1 | 1 | 1 | 1 | 1 | 1 |  | - | - | 1 | 1 | 1 | 1 | 1 | 1 | 1 | 1 |  |  |
| Gait | - | - | - | - | - | - | - | - | - | - | - |  | - | - | - | - | - | - | - | - | - | - |  |  |
| Palpebral Size | - | - | - | - | - | - | - | - | - | - | - |  | - | - | - | - | - | - | - | - | - | - |  |  |
| Lacrimation | - | - | - | - | - | - | - | - | - | - | - |  | - | - | - | - | - | - | - | - | - | - |  |  |
| Skin Color | - | - | - | - | - | - | - | - | - | - | - |  | - | - | - | - | - | - | - | - | - | - |  |  |
| Salivation | - | - | - | - | - | - | - | - | - | - | - |  | - | - | - | - | - | - | - | - | - | - |  |  |
| Diarrhea | - | - | - | - | - | - | - | - | - | - | - |  | - | - | - | - | - | - | - | - | - | - |  |  |
| Convulsion/Tremors | - | - | - | - | - | - | - | - | - | - | - |  | - | - | - | - | - | - | - | - | - | - |  |  |
| Others | - | - | - | - | - | - | - | - | - | - | - |  | - | - | - | - | - | - | - | - | - | - |  |  |
| Death | - | - | - | - | - | - | - | - | - | - | - |  | - | - | - | - | - | - | - | - | - | - |  |  |

**Notes (alphabetical order):**

**Table 3-17.** MTD results of ISFP10, cage side observation at the 2^nd^ day after the last dosing.

| Gender | Female |  |  |  |  |  |  |  |  |  |  |  | Male |  |  |  |  |  |  |  |  |  |  |  |
| --- | --- | --- | --- | --- | --- | --- | --- | --- | --- | --- | --- | --- | --- | --- | --- | --- | --- | --- | --- | --- | --- | --- | --- | --- |
| Test Article | Vehicle |  | PT# 1288839<br>(NID-490)<br>(ISFP10) |  |  |  |  |  |  |  |  |  | Vehicle |  | PT# 1288839<br>(NID-490)<br>(ISFP10) |  |  |  |  |  |  |  |  |  |
| Route/Schedule | IP, BID,<br>q12h x 7<br>days |  | IP, BID, q12h<br>x 7 days |  |  |  |  |  |  |  |  |  | IP, BID,<br>q12h x 7<br>days |  | IP, BID, q12h<br>x 7 days |  |  |  |  |  |  |  |  |  |
| Dosage | 10<br>mL/kg |  | 12.5<br>mg/kg |  | 25<br>mg/kg |  | 50<br>mg/kg |  | 100<br>mg/kg |  | 200<br>mg/kg |  | 10<br>mL/kg |  | 12.5<br>mg/kg |  | 25<br>mg/kg |  | 50<br>mg/kg |  | 100<br>mg/kg |  | 200<br>mg/kg |  |
| BEHAVIORAL | #1 | #2 | #1 | #2 | #1 | #2 | #1 | #2 | #1 | #2 | #1 | #2 | #1 | #2 | #1 | #2 | #1 | #2 | #1 | #2 | #1 | #2 | #1 | #2 |
| Body weight (g) | 21 | 20 | 20 | 20 | 21 | 20 | 20 | 21 | 20 | 19 | 19 |  | 23 | 25 | 23 | 24 | 22 | 23 | 23 | 24 | 22 | 21 |  |  |
| Body temperature | - | - | - | - | - | - | - | - | - | - | - |  | - | - | - | - | - | - | - | - | - | - |  |  |
| Respiration | - | - | - | - | - | - | - | - | - | - | - |  | - | - | - | - | - | - | - | - | - | - |  |  |
| Hunched back | - | - | - | - | - | - | ± | ± | ± | ± | ± |  | - | - | ± | ± | ± | ± | ± | ± | ± | ± |  |  |
| Piloerection | - | - | - | - | - | ± | ± | ± | ± | ± | ± |  | - | - | ± | ± | ± | ± | ± | ± | ± | ± |  |  |
| Activity | 1 | 1 | 1 | 1 | 1 | 1 | 1 | 1 | 1 | 1 | 1 |  | - | - | 1 | 1 | 1 | 1 | 1 | 1 | 1 | 1 |  |  |
| Gait | - | - | - | - | - | - | - | - | - | - | - |  | - | - | - | - | - | - | - | - | - | - |  |  |
| Palpebral Size | - | - | - | - | - | - | - | - | - | - | - |  | - | - | - | - | - | - | - | - | - | - |  |  |
| Lacrimation | - | - | - | - | - | - | - | - | - | - | - |  | - | - | - | - | - | - | - | - | - | - |  |  |
| Skin Color | - | - | - | - | - | - | - | - | - | - | - |  | - | - | - | - | - | - | - | - | - | - |  |  |
| Salivation | - | - | - | - | - | - | - | - | - | - | - |  | - | - | - | - | - | - | - | - | - | - |  |  |
| Diarrhea | - | - | - | - | - | - | - | - | - | - | - |  | - | - | - | - | - | - | - | - | - | - |  |  |
| Convulsion/Tremors | - | - | - | - | - | - | - | - | - | - | - |  | - | - | - | - | - | - | - | - | - | - |  |  |
| Others | - | - | - | - | - | - | - | - | - | - | - |  | - | - | - | - | - | - | - | - | - | - |  |  |
| Death | - | - | - | - | - | - | - | - | - | - | - |  | - | - | - | - | - | - | - | - | - | - |  |  |

**Table 3-18.** MTD results of ISFP10, cage side observation at the 3^rd^ day after the last dosing.

| Gender | Female |  |  |  |  |  |  |  |  |  |  |  | Male |  |  |  |  |  |  |  |  |  |  |  |
| --- | --- | --- | --- | --- | --- | --- | --- | --- | --- | --- | --- | --- | --- | --- | --- | --- | --- | --- | --- | --- | --- | --- | --- | --- |
| Test Article | Vehicle |  | PT# 1288839<br>(NID-490)<br>(ISFP10) |  |  |  |  |  |  |  |  |  | Vehicle |  | PT# 1288839<br>(NID-490)<br>(ISFP10) |  |  |  |  |  |  |  |  |  |
| Route/Schedule | IP, BID,<br>q12h x 7<br>days |  | IP, BID, q12h<br>x 7 days |  |  |  |  |  |  |  |  |  | IP, BID,<br>q12h x 7<br>days |  | IP, BID, q12h<br>x 7 days |  |  |  |  |  |  |  |  |  |
| Dosage | 10<br>mL/kg |  | 12.5<br>mg/kg |  | 25<br>mg/kg |  | 50<br>mg/kg |  | 100<br>mg/kg |  | 200<br>mg/kg |  | 10<br>mL/kg |  | 12.5<br>mg/kg |  | 25<br>mg/kg |  | 50<br>mg/kg |  | 100<br>mg/kg |  | 200<br>mg/kg |  |
| BEHAVIORAL | #1 | #2 | #1 | #2 | #1 | #2 | #1 | #2 | #1 | #2 | #1 | #2 | #1 | #2 | #1 | #2 | #1 | #2 | #1 | #2 | #1 | #2 | #1 | #2 |
| Body weight (g) | 21 | 20 | 20 | 20 | 22 | 20 | 21 | 20 | 20 | 20 | 20 |  | 24 | 25 | 24 | 24 | 23 | 23 | 25 | 23 | 24 | 22 |  |  |
| Body temperature | - | - | - | - | - | - | - | - | - | - | - |  | - | - | - | - | - | - | - | - | - | - |  |  |
| Respiration | - | - | - | - | - | - | - | - | - | - | - |  | - | - | - | - | - | - | - | - | - | - |  |  |
| Hunched back | - | - | - | - | - | - | ± | ± | ± | ± | ± |  | - | - | ± | ± | ± | ± | ± | ± | ± | ± |  |  |
| Piloerection | - | - | - | - | - | - | ± | ± | ± | ± | ± |  | - | - | ± | ± | ± | ± | ± | ± | ± | ± |  |  |
| Activity | 1 | 1 | 1 | 1 | 1 | 1 | 1 | 1 | 1 | 1 | 1 |  | - | - | 1 | 1 | 1 | 1 | 1 | 1 | 1 | 1 |  |  |
| Gait | - | - | - | - | - | - | - | - | - | - | - |  | - | - | - | - | - | - | - | - | - | - |  |  |
| Palpebral Size | - | - | - | - | - | - | - | - | - | - | - |  | - | - | - | - | - | - | - | - | - | - |  |  |
| Lacrimation | - | - | - | - | - | - | - | - | - | - | - |  | - | - | - | - | - | - | - | - | - | - |  |  |
| Skin Color | - | - | - | - | - | - | - | - | - | - | - |  | - | - | - | - | - | - | - | - | - | - |  |  |
| Salivation | - | - | - | - | - | - | - | - | - | - | - |  | - | - | - | - | - | - | - | - | - | - |  |  |
| Diarrhea | - | - | - | - | - | - | - | - | - | - | - |  | - | - | - | - | - | - | - | - | - | - |  |  |
| Convulsion/Tremors | - | - | - | - | - | - | - | - | - | - | - |  | - | - | - | - | - | - | - | - | - | - |  |  |
| Others | - | - | - | - | - | - | - | - | - | - | - |  | - | - | - | - | - | - | - | - | - | - |  |  |
| Death | - | - | - | - | - | - | - | - | - | - | - |  | - | - | - | - | - | - | - | - | - | - |  |  |

**Notes (alphabetical order):**

**Table 4-1.** MTD results, body weight records post dose administrations, female.

| Group | Test Article | Dose Route/<br>Schedule | Animal<br>No. | Body Weight (g) |  |  |  |  |  |  |  |  |  |  |  |  |  |  |  |  |
| --- | --- | --- | --- | --- | --- | --- | --- | --- | --- | --- | --- | --- | --- | --- | --- | --- | --- | --- | --- | --- |
|  |  |  |  | Day 0 |  | Day 1 |  | Day 2 |  | Day 3 |  | Day 4 |  | Day 5 |  | Day 6 |  | Day 7 | Day 8 | Day 9 |
|  |  |  |  | 1<sup>st</sup><br>dose | 2<sup>nd</sup><br>dose | 3<sup>rd</sup><br>dose | 4<sup>th</sup><br>dose | 5<sup>th</sup><br>dose | 6<sup>th</sup><br>dose | 7<sup>th</sup><br>dose | 8<sup>th</sup><br>dose | 9<sup>th</sup><br>dose | 10<sup>th</sup><br>dose | 11<sup>th</sup><br>dose | 12<sup>th</sup><br>dose | 13<sup>th</sup><br>dose | 14<sup>th</sup><br>dose | 1<sup>st</sup> day<br>after<br>last<br>dose | 2<sup>nd</sup> day<br>after<br>last<br>dose | 3<sup>rd</sup> day<br>after<br>last<br>dose |
| 1 | Vehicle | 10 mL/kg<br>IP, BID,<br>q12h<br>x 7 days | 1 | 20 | 19 | 20 | 19 | 21 | 20 | 22 | 21 | 22 | 21 | 21 | 20 | 22 | 20 | 22 | 21 | 21 |
|  |  |  | 2 | 18 | 18 | 19 | 18 | 19 | 19 | 20 | 19 | 20 | 19 | 20 | 19 | 20 | 19 | 20 | 20 | 20 |
|  |  |  | Mean | 19.0 | 18.5 | 19.5 | 18.5 | 20.0 | 19.5 | 21.0 | 20.0 | 21.0 | 20.0 | 20.5 | 19.5 | 21.0 | 19.5 | 21.0 | 20.5 | 20.5 |
|  |  |  | SEM | 1.0 | 0.5 | 0.5 | 0.5 | 1.0 | 0.5 | 1.0 | 1.0 | 1.0 | 1.0 | 0.5 | 0.5 | 1.0 | 0.5 | 1.0 | 0.5 | 0.5 |
| 2 | PT# 1288839<br>(NID-490)<br>(ISFP10) | 12.5 mg/kg<br>IP, BID,<br>q12h<br>x 7 days | 1 | 19 | 19 | 19 | 18 | 19 | 19 | 19 | 19 | 20 | 20 | 20 | 19 | 20 | 19 | 20 | 20 | 20 |
|  |  |  | 2 | 20 | 19 | 19 | 18 | 19 | 19 | 19 | 19 | 19 | 19 | 19 | 20 | 19 | 20 | 21 | 20 | 20 |
|  |  |  | Mean | 19.5 | 19.0 | 19.0 | 18.0 | 19.0 | 19.0 | 19.0 | 19.0 | 19.5 | 19.5 | 19.5 | 19.5 | 19.5 | 19.5 | 20.5 | 20.0 | 20.0 |
|  |  |  | SEM | 0.5 | 0.0 | 0.0 | 0.0 | 0.0 | 0.0 | 0.0 | 0.0 | 0.5 | 0.5 | 0.5 | 0.5 | 0.5 | 0.5 | 0.5 | 0.0 | 0.0 |
| 3 | PT# 1288839<br>(NID-490)<br>(ISFP10) | 25 mg/kg<br>IP BID, q12h<br>x 7 days | 1 | 21 | 20 | 19 | 19 | 20 | 20 | 21 | 20 | 21 | 20 | 20 | 20 | 21 | 20 | 21 | 21 | 22 |
|  |  |  | 2 | 19 | 18 | 18 | 17 | 17 | 17 | 18 | 17 | 19 | 18 | 18 | 18 | 19 | 19 | 20 | 20 | 20 |
|  |  |  | Mean | 20.0 | 19.0 | 18.5 | 18.0 | 18.5 | 18.5 | 19.5 | 18.5 | 20.0 | 19.0 | 19.0 | 19.0 | 20.0 | 19.5 | 20.5 | 20.5 | 21.0 |
|  |  |  | SEM | 1.0 | 1.0 | 0.5 | 1.0 | 1.5 | 1.5 | 1.5 | 1.5 | 1.0 | 1.0 | 1.0 | 1.0 | 1.0 | 0.5 | 0.5 | 0.5 | 1.0 |
| 4 | PT# 1288839<br>(NID-490)<br>(ISFP10) | 50 mg/kg<br>IP, BID,<br>q12h<br>x 7 days | 1 | 20 | 20 | 19 | 18 | 18 | 19 | 20 | 19 | 20 | 19 | 20 | 20 | 21 | 20 | 20 | 20 | 21 |
|  |  |  | 2 | 20 | 19 | 19 | 18 | 19 | 18 | 20 | 19 | 20 | 19 | 19 | 19 | 18 | 19 | 19 | 21 | 20 |
|  |  |  | Mean | 20.0 | 19.5 | 19.0 | 18.0 | 18.5 | 18.5 | 20.0 | 19.0 | 20.0 | 19.0 | 19.5 | 19.5 | 19.5 | 19.5 | 19.5 | 20.5 | 20.5 |
|  |  |  | SEM | 0.0 | 0.5 | 0.0 | 0.0 | 0.5 | 0.5 | 0.0 | 0.0 | 0.0 | 0.0 | 0.5 | 0.5 | 1.5 | 0.5 | 0.5 | 0.5 | 0.5 |
| 5 | PT# 1288839<br>(NID-490)<br>(ISFP10) | 100 mg/kg<br>IP, BID,<br>q12h<br>x 7 days | 1 | 19 | 19 | 19 | 19 | 20 | 19 | 20 | 20 | 21 | 20 | 21 | 20 | 20 | 19 | 19 | 20 | 20 |
|  |  |  | 2 | 20 | 19 | 19 | 20 | 20 | 19 | 20 | 19 | 19 | 19 | 19 | 19 | 20 | 19 | 20 | 19 | 20 |
|  |  |  | Mean | 19.5 | 19.0 | 19.0 | 19.5 | 20.0 | 19.0 | 20.0 | 19.5 | 20.0 | 19.5 | 20.0 | 19.5 | 20.0 | 19.0 | 19.5 | 19.5 | 20.0 |
|  |  |  | SEM | 0.5 | 0.0 | 0.0 | 0.5 | 0.0 | 0.0 | 0.0 | 0.5 | 1.0 | 0.5 | 1.0 | 0.5 | 0.0 | 0.0 | 0.5 | 0.5 | 0.0 |
| 6 | PT# 1288839<br>(NID-490)<br>(ISFP10) | 200 mg/kg<br>IP, BID,<br>q12h<br>x 7 days | 1 | 21 | 20 | 19 | 19 | 18 | 18 | 18 | 18 | 18 | 18 | 18 | 19 | 19 | 19 | 19 | 19 | 20 |
|  |  |  | 2 | 21 | 19 | 18 | 18 | 18 | 17 | 17 | HS |  |  |  |  |  |  |  |  |  |
|  |  |  | Mean | 21.0 | 19.5 | 18.5 | 18.5 | 18.0 | 17.5 | 17.5 | 18.0 | 18.0 | 18.0 | 18.0 | 19.0 | 19.0 | 19.0 | 19.0 | 19.0 | 20.0 |
|  |  |  | SEM | 0.0 | 0.5 | 0.5 | 0.5 | 0.0 | 0.5 | 0.5 | -- | -- | -- | -- | -- | -- | -- | -- | -- | -- |

**Table 4-2.** MTD results, body weight records post dose administrations, male.

| Group | Test Article | Dose Route/<br>Schedule | Animal<br>No. | Body Weight (g) |  |  |  |  |  |  |  |  |  |  |  |  |  |  |  |  |
| --- | --- | --- | --- | --- | --- | --- | --- | --- | --- | --- | --- | --- | --- | --- | --- | --- | --- | --- | --- | --- |
|  |  |  |  | Day 0 |  | Day 1 |  | Day 2 |  | Day 3 |  | Day 4 |  | Day 5 |  | Day 6 |  | Day 7 | Day 8 | Day 9 |
|  |  |  |  | 1<sup>st</sup><br>dose | 2<sup>nd</sup><br>dose | 3<sup>rd</sup><br>dose | 4<sup>th</sup><br>dose | 5<sup>th</sup><br>dose | 6<sup>th</sup><br>dose | 7<sup>th</sup><br>dose | 8<sup>th</sup><br>dose | 9<sup>th</sup><br>dose | 10<sup>th</sup><br>dose | 11<sup>th</sup><br>dose | 12<sup>th</sup><br>dose | 13<sup>th</sup><br>dose | 14<sup>th</sup><br>dose | 1<sup>st</sup> day<br>after<br>last<br>dose | 2<sup>nd</sup> day<br>after<br>last<br>dose | 3<sup>rd</sup> day<br>after<br>last<br>dose |
| 1 | Vehicle | 10 mL/kg<br>IP, BID,<br>q12h<br>x 7 days | 1 | 23 | 23 | 24 | 23 | 24 | 24 | 24 | 24 | 25 | 23 | 24 | 23 | 24 | 23 | 24 | 23 | 24 |
|  |  |  | 2 | 24 | 22 | 22 | 21 | 22 | 22 | 23 | 22 | 23 | 22 | 23 | 22 | 24 | 24 | 25 | 25 | 25 |
|  |  |  | Mean | 23.5 | 22.5 | 23.0 | 22.0 | 23.0 | 23.0 | 23.5 | 23.0 | 24.0 | 22.5 | 23.5 | 22.5 | 24.0 | 23.5 | 24.5 | 24.0 | 24.5 |
|  |  |  | SEM | 0.5 | 0.5 | 1.0 | 1.0 | 1.0 | 1.0 | 0.5 | 1.0 | 1.0 | 0.5 | 0.5 | 0.5 | 0.0 | 0.5 | 0.5 | 1.0 | 0.5 |
| 2 | PT# 1288839<br>(NID-490)<br>(ISFP10) | 12.5 mg/kg<br>IP, BID,<br>q12h<br>x 7 days | 1 | 24 | 22 | 23 | 22 | 23 | 22 | 24 | 22 | 23 | 21 | 23 | 22 | 23 | 22 | 24 | 23 | 24 |
|  |  |  | 2 | 23 | 22 | 23 | 22 | 23 | 23 | 24 | 23 | 24 | 23 | 24 | 23 | 24 | 23 | 24 | 24 | 24 |
|  |  |  | Mean | 23.5 | 22.0 | 23.0 | 22.0 | 23.0 | 22.5 | 24.0 | 22.5 | 23.5 | 22.0 | 23.5 | 22.5 | 23.5 | 22.5 | 24.0 | 23.5 | 24.0 |
|  |  |  | SEM | 0.5 | 0.0 | 0.0 | 0.0 | 0.0 | 0.5 | 0.0 | 0.5 | 0.5 | 1.0 | 0.5 | 0.5 | 0.5 | 0.5 | 0.0 | 0.5 | 0.0 |
| 3 | PT# 1288839<br>(NID-490)<br>(ISFP10) | 25 mg/kg<br>IP BID, q12h<br>x 7 days | 1 | 23 | 22 | 22 | 21 | 22 | 21 | 23 | 22 | 23 | 22 | 24 | 22 | 22 | 22 | 23 | 22 | 23 |
|  |  |  | 2 | 22 | 21 | 21 | 21 | 22 | 21 | 22 | 22 | 23 | 21 | 23 | 21 | 21 | 22 | 23 | 23 | 23 |
|  |  |  | Mean | 22.5 | 21.5 | 21.5 | 21.0 | 22.0 | 21.0 | 22.5 | 22.0 | 23.0 | 21.5 | 23.5 | 21.5 | 21.5 | 22.0 | 23.0 | 22.5 | 23.0 |
|  |  |  | SEM | 0.5 | 0.5 | 0.5 | 0.0 | 0.0 | 0.0 | 0.5 | 0.0 | 0.0 | 0.5 | 0.5 | 0.5 | 0.5 | 0.0 | 0.0 | 0.5 | 0.0 |
| 4 | PT# 1288839<br>(NID-490)<br>(ISFP10) | 50 mg/kg<br>IP, BID,<br>q12h<br>x 7 days | 1 | 23 | 21 | 21 | 21 | 22 | 21 | 23 | 21 | 22 | 21 | 22 | 21 | 23 | 21 | 23 | 23 | 25 |
|  |  |  | 2 | 24 | 23 | 23 | 22 | 23 | 22 | 23 | 22 | 23 | 22 | 22 | 22 | 23 | 22 | 23 | 24 | 23 |
|  |  |  | Mean | 23.5 | 22.0 | 22.0 | 21.5 | 22.5 | 21.5 | 23.0 | 21.5 | 22.5 | 21.5 | 22.0 | 21.5 | 23.0 | 21.5 | 23.0 | 23.5 | 24.0 |
|  |  |  | SEM | 0.5 | 1.0 | 1.0 | 0.5 | 0.5 | 0.5 | 0.0 | 0.5 | 0.5 | 0.5 | 0.0 | 0.5 | 0.0 | 0.5 | 0.0 | 0.5 | 1.0 |
| 5 | PT# 1288839<br>(NID-490)<br>(ISFP10) | 100 mg/kg<br>IP, BID,<br>q12h<br>x 7 days | 1 | 20 | 22 | 22 | 22 | 22 | 22 | 21 | 22 | 22 | 21 | 22 | 21 | 23 | 22 | 22 | 22 | 24 |
|  |  |  | 2 | 22 | 21 | 21 | 20 | 21 | 20 | 20 | 20 | 21 | 20 | 21 | 21 | 20 | 20 | 21 | 21 | 22 |
|  |  |  | Mean | 21.0 | 21.5 | 21.5 | 21.0 | 21.5 | 21.0 | 20.5 | 21.0 | 21.5 | 20.5 | 21.5 | 21.0 | 21.5 | 21.0 | 21.5 | 21.5 | 23.0 |
|  |  |  | SEM | 1.0 | 0.5 | 0.5 | 1.0 | 0.5 | 1.0 | 0.5 | 1.0 | 0.5 | 0.5 | 0.5 | 0.0 | 1.5 | 1.0 | 0.5 | 0.5 | 1.0 |
| 6 | PT# 1288839<br>(NID-490)<br>(ISFP10) | 200 mg/kg<br>IP, BID,<br>q12h<br>x 7 days | 1 | 22 | 21 | 21 | 21 | Dead |  |  |  |  |  |  |  |  |  |  |  |  |
|  |  |  | 2 | 21 | 20 | 19 | 19 | Dead |  |  |  |  |  |  |  |  |  |  |  |  |
|  |  |  | Mean | 21.5 | 20.5 | 20.0 | 20.0 | -- | -- | -- | -- | -- | -- | -- | -- | -- | -- | -- | -- | -- |
|  |  |  | SEM | 0.5 | 0.5 | 1.0 | 1.0 |  |  |  |  |  |  |  |  |  |  |  |  |  |

**Table 5-1.**
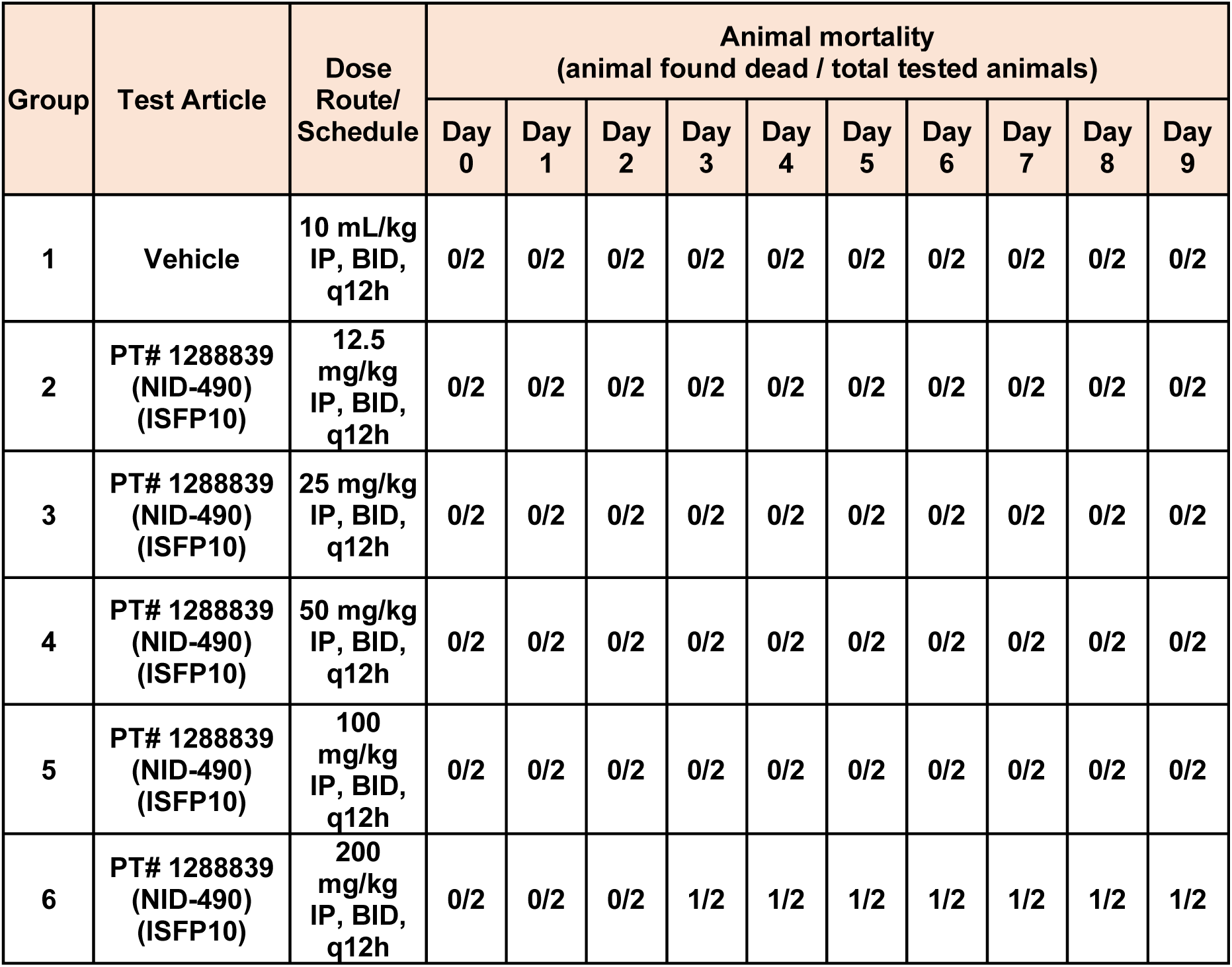
MTD results, mortality assessment prior and post dose administrations, female.

**Table 5-2.** MTD results, mortality assessment prior and post dose administrations, male.

| Group | Test Article | Dose Route/<br>Schedule | Animal mortality<br>(animal found dead / total tested animals) |  |  |  |  |  |  |  |  |  |
| --- | --- | --- | --- | --- | --- | --- | --- | --- | --- | --- | --- | --- |
|  |  |  | Day 0 | Day 1 | Day 2 | Day 3 | Day 4 | Day 5 | Day 6 | Day 7 | Day 8 | Day 9 |
| 1 | Vehicle | 10 mL/kg<br>IP, BID,<br>q12h | 0/2 | 0/2 | 0/2 | 0/2 | 0/2 | 0/2 | 0/2 | 0/2 | 0/2 | 0/2 |
| 2 | PT# 1288839<br>(NID-490)<br>(ISFP10) | 12.5<br>mg/kg<br>IP, BID,<br>q12h | 0/2 | 0/2 | 0/2 | 0/2 | 0/2 | 0/2 | 0/2 | 0/2 | 0/2 | 0/2 |
| 3 | PT# 1288839<br>(NID-490)<br>(ISFP10) | 25 mg/kg<br>IP, BID,<br>q12h | 0/2 | 0/2 | 0/2 | 0/2 | 0/2 | 0/2 | 0/2 | 0/2 | 0/2 | 0/2 |
| 4 | PT# 1288839<br>(NID-490)<br>(ISFP10) | 50 mg/kg<br>IP, BID,<br>q12h | 0/2 | 0/2 | 0/2 | 0/2 | 0/2 | 0/2 | 0/2 | 0/2 | 0/2 | 0/2 |
| 5 | PT# 1288839<br>(NID-490)<br>(ISFP10) | 100<br>mg/kg<br>IP, BID,<br>q12h | 0/2 | 0/2 | 0/2 | 0/2 | 0/2 | 0/2 | 0/2 | 0/2 | 0/2 | 0/2 |
| 6 | PT# 1288839<br>(NID-490)<br>(ISFP10) | 200<br>mg/kg<br>IP, BID,<br>q12h | 0/2 | 0/2 | 2/2 | 2/2 | 2/2 | 2/2 | 2/2 | 2/2 | 2/2 | 2/2 |

#### Female mice

200 mg/kg dose groups exhibited mild to moderate reductions in abdominal muscle tone. Beginning with the third dose, signs of mild to moderate piloerection, reduced palpebral aperture, and a hunched posture were observed in the 25, 50, 100, and 200 mg/kg groups. One animal in the 200 mg/kg group was humanely euthanized 30 minutes after the eighth dose (Day 3) due to severe adverse effects noted following the seventh administration. These included hypothermia, bradypnea, hunched posture, piloerection, markedly reduced activity (unresponsive to stimulation), decreased palpebral size, wet tail, and a moribund state. At the lowest dose of 12.5 mg/kg, only one animal exhibited transient mild to moderate piloerection and hunched posture, observed 30 minutes after the eleventh IP dose (**Table 3-12**). Apart from the euthanized animal in the 200 mg/kg group, no significant body weight loss was detected across any dose groups (**Table 4-1**).

In summary, ISFP10 administered at 200 mg/kg IP BID for three days resulted in 50% mortality and was deemed intolerable. In contrast, dosing at 12.5, 25, 50, and 100 mg/kg BID for seven days was well tolerated, with no mortality observed (**Table 5-1**).

#### Male mice

On Day 0, within 30 minutes of the first intraperitoneal (IP) dose, only animals in the 100 and 200 mg/kg groups exhibited mild to moderate reductions in abdominal and limb tone. Beginning with the second dose, animals in the 200 mg/kg group developed progressive adverse symptoms, including mild to severe lacrimation, hypothermia, bradypnea, hunched posture, piloerection, markedly reduced activity (no ambulation even upon stimulation), decreased palpebral aperture, ocular discharge, and wet tail (**Table 3-3**). These clinical signs worsened over time, and both animals in this group were found deceased between the fourth and fifth doses (Day 1 and Day 2, respectively). At lower dose levels (12.5, 25, 50, and 100 mg/kg), animals exhibited mild to moderate adverse effects—such as piloerection, decreased palpebral size, hunched posture, and wet tail—but no severe toxicity was observed. Body weight remained stable in all surviving animals (**Table 4-2**).

In summary, ISFP10 administered at 200 mg/kg BID q12h resulted in 100% mortality within two days and was deemed intolerable. In contrast, repeated IP dosing at 12.5, 25, 50, and 100 mg/kg BID for seven consecutive days was well tolerated, with no mortality observed (**Table 5-2**).

#### Plasma PK for ISFP10

The concentration-time profile of ISFP10 in plasma following IP BID × 7 days administrations of ISFP10 is plotted in Figure **1A**. The PK parameters, including *t*_1/2_, *T*_max_, *C*_max_, AUC_last_, AUC_Inf_, AUC/D, AUCExtr, MRT, Vz_F and CL_F, are provided in **Table 6-1** and were calculated using noncompartmental analysis (NCA) with WinNonlin. The ISFP10 dosing at 12.5, 25, 50 and 100 mg/kg IP BID for 7 days yielded plasma C_max_ values that ranged from 3.450 to 5.140 µg/mL b, R^2^ = 0.6916. The respective AUC_last_ values increased with dose, ranging from 30.463 to 81.717 h*µg/mL with moderate linearity, R^2^ = 0.8602

**Fig. 1.**
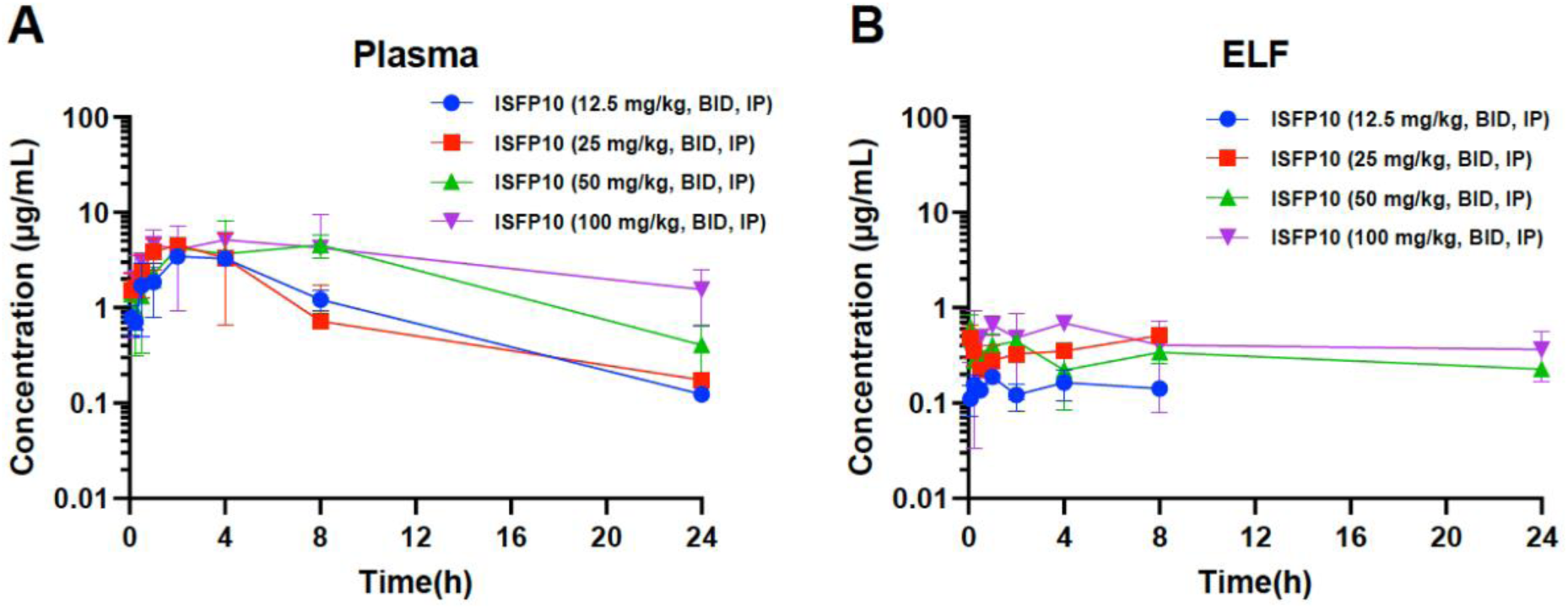
The mean plasma (A) and ELF (B) concentration-time profiles of ISFP10 IP BID × 7 days administrations to uninfected C57BL/6 mice. Three animals were sacrificed per time point at 8 time points, 0.083, 0.25, 0.5, 1, 2, 4, 8, and 24 h, after the last intraperitoneal (IP) administration. Error bars are SD values.

**Table 6-1.** The plasma PK parameters of ISFP10 by IP twice daily administrations for 7 days to uninfected C57BL/6 mice with Best fit.

| Dose, IP<br>(mg/kg) | Time<br>point<br>(h) | t<sub>1/2</sub><br>(h) | T<sub>max</sub><br>(h) | C<sub>max</sub><br>(µg/mL) | AUC<sub>last</sub><br>(h*µg/mL) | AUC<sub>Inf</sub><br>(h*µg/mL) | AUC/D<br>(h*kg*µg/mL/mg) | AUC<br>Extr<br>(%) | MRT<br>(h) | Vz_F<br>(L/kg) | CL_F<br>(mL/min/kg) |
| --- | --- | --- | --- | --- | --- | --- | --- | --- | --- | --- | --- |
| 12.5 | 4.0-24.0 | 4.39 | 2.00 | 3.450 | 30.463 | 31.249 | 1.25 | 2.51 | 6.29 | 5.07 | 13.33 |
| 25 | 4.0-24.0 | 5.31 | 2.00 | 4.588 | 29.793 | 31.134 | 0.62 | 4.31 | 6.13 | 12.31 | 26.77 |
| 50 | NA | NA | 8.00 | 4.536 | 68.525 | NA | NA | NA | NA | NA | NA |
| 100 | NA | NA | 4.00 | 5.140 | 81.717 | NA | NA | NA | NA | NA | NA |

#### Epithelial Ling Fluid Pharmacokinetics for ISFP10

The concentration-time profile of ISFP10 in epithelial lining fluid (ELF) following IP administrations of ISFP10 is plotted in Figure **1B** using the information tabulated in **Table 6-2**. ISFP10 at 12.5, 25, 50 and 100 mg/kg IP BID × 7 days, yielded concentrations in ELF in the 100 mg/kg dose group that ranged from 0.345 – 0.690 µg/mL over the 24 h period. The 12.5, 25, and 50 mg/kg dose groups yielded measurable concentrations up to 8 h post dose with respective ranges of 0.111 – 0.189; 0.219 – 0.509; and 0.221 – 0.617 µg/mL that fluctuated over time.

**Table 6-2.** The ELF PK parameters of ISFP10 by IP twice daily administrations for 7 days to uninfected C57BL/6 mice with Best fit.

| Dose,<br>IP<br>(mg/kg) | Time<br>point<br>(h) | t<sub>1/2</sub><br>(h) | T<sub>max</sub><br>(h) | C<sub>max</sub><br>(µg/mL) | AUC<sub>last</sub><br>(h*µg/mL) | AUC<sub>Inf</sub><br>(h*µg/mL) | AUC/D<br>(h*kg*µg/mL/mg) | AUC<br>Extr<br>(%) | MRT<br>(h) | Vz_F<br>(L/kg) | CL_F<br>(mL/min/kg) | ELF / Plasma |  |
| --- | --- | --- | --- | --- | --- | --- | --- | --- | --- | --- | --- | --- | --- |
|  |  |  |  |  |  |  |  |  |  |  |  | C<sub>max</sub> | AUC<sub>last</sub> |
| 12.5 | NA | NA | 1.000 | 0.189 | 1.197 | NA | NA | NA | NA | NA | NA | 5.48% | 3.93% |
| 25 | NA | NA | 8.000 | 0.509 | 2.991 | NA | NA | NA | NA | NA | NA | 11.09% | 10.04% |
| 50 | 0.5-24 | 35.66 | 0.083 | 0.617 | 7.103 | 18.782 | 0.19 | 62.18 | 50.92 | 273.93 | 88.74 | 13.60% | 10.37% |
| 100 | NA | NA | 4.000 | 0.690 | 10.561 | NA | NA | NA | NA | NA | NA | 13.42% | 12.92% |

The ISFP10 dosing of 12.5, 25, 50 and 100 mg/kg IP BID × 7 days yielded C_max_ values that ranged from 0.189 to 0.690 µg/mL and the AUC_last_ values that ranged from 1.197 to 10.561 h*µg/mL (**Table 6-2**). The ELF penetration values (AUC_last_ ELF/AUC_last_ plasma) ranged from 3.93% to 12.92% with the tested doses of 12.5, 25, 50 and 100 mg/kg IP BID × 7 days (**Figure 2**). The ELF AUC_last_ parameters correlated linearly with dose with an R-squared value of 0.9546. The ELF C_max_ values did not correlate linearly with dose, R-squared value of 0.6667 (**Figures 3** and **4**).

**Fig. 2.**
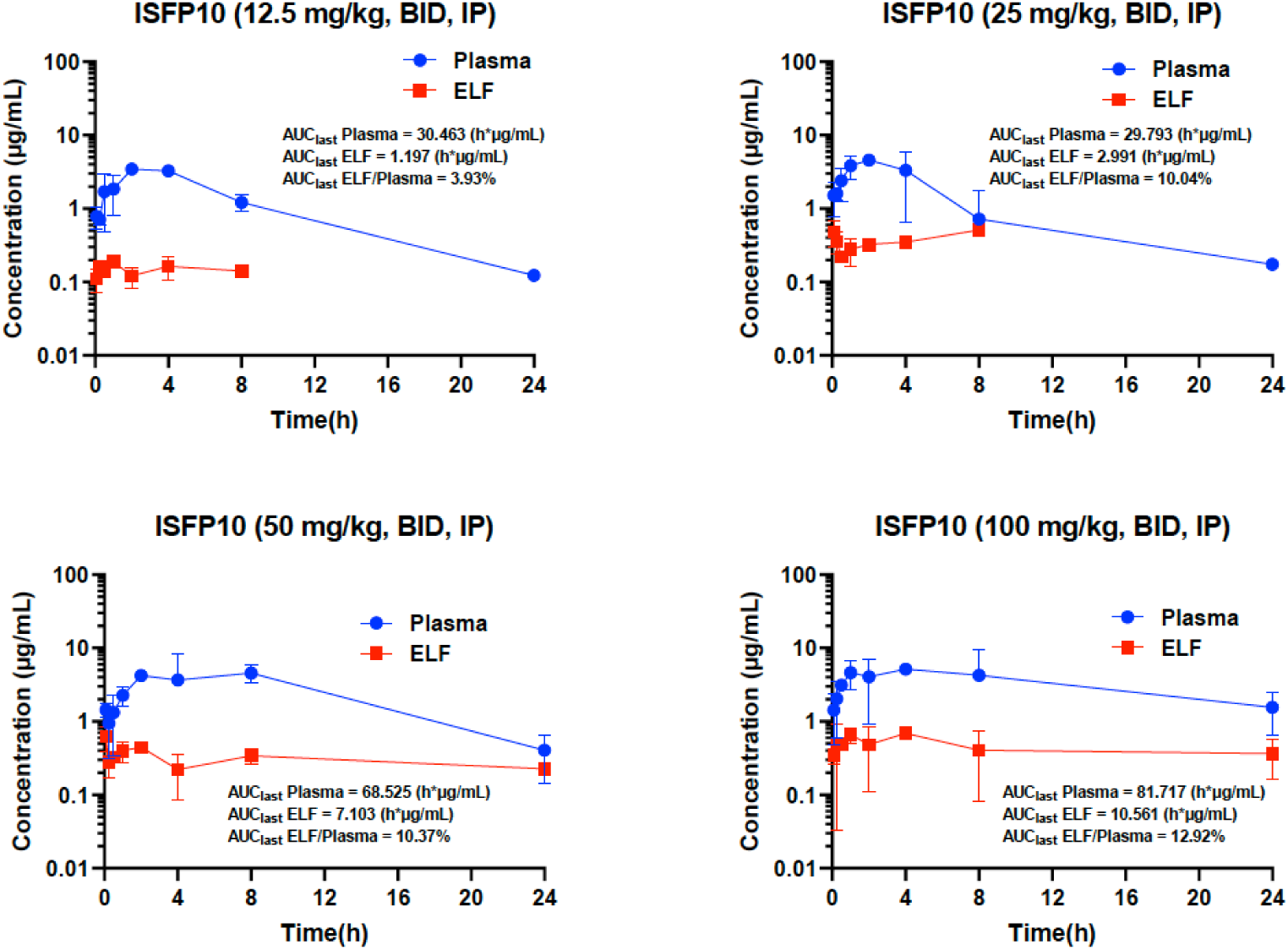
The mean ELF (red): plasma (blue) concentration ratio over time profiles of ISFP10 IP BID × 7 days administrations to uninfected C57BL/6 mice. Three animals were sacrificed per time point at 8 time points, 0.083, 0.25, 0.5, 1, 2, 4, 8, and 24 h, after the last intraperitoneal (IP) administration. Error bars are SD values.

**Fig. 3.**
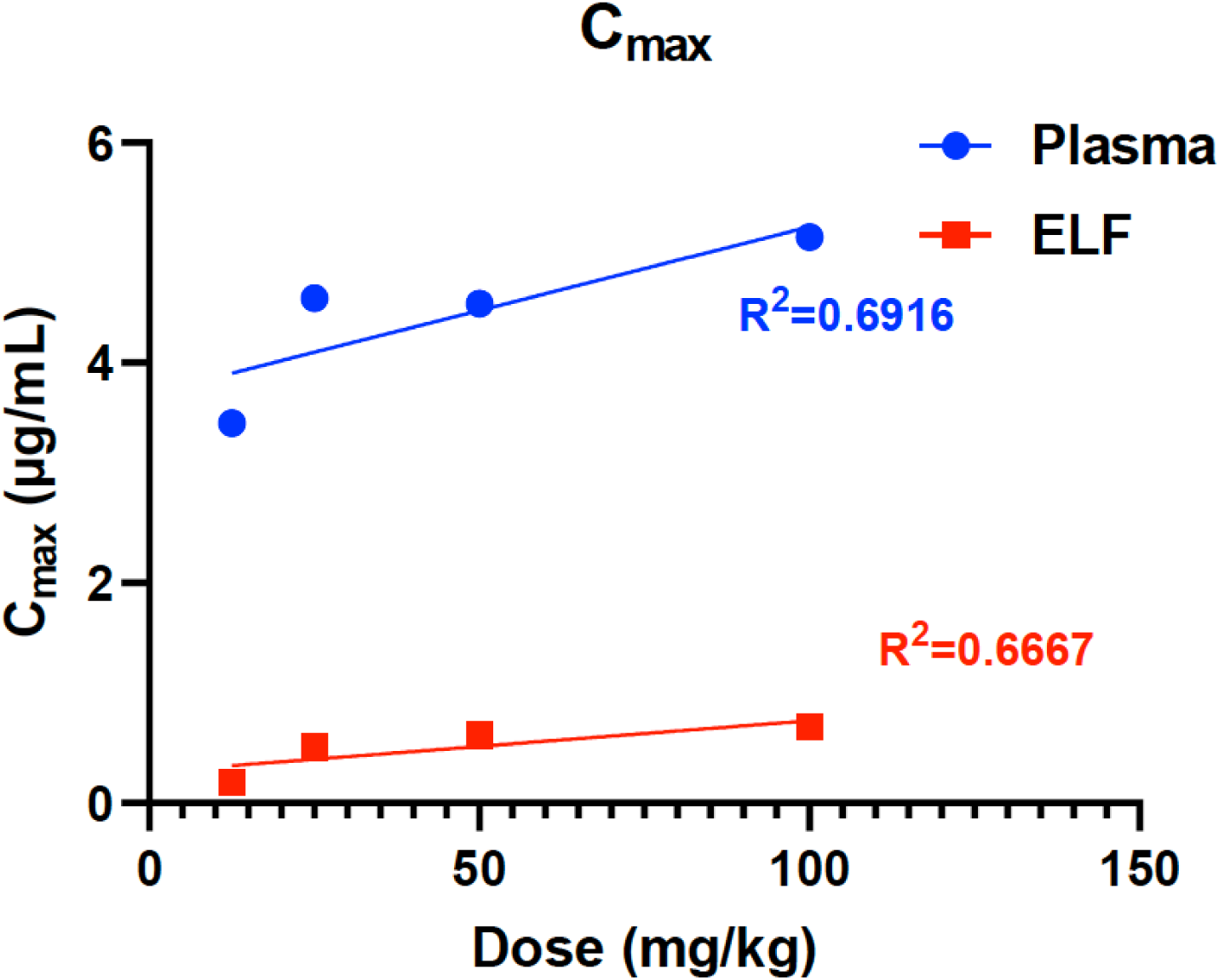
Dose correlation of ISFP10 at IP administration with C_max_ in plasma and ELF (µg/mL) of uninfected C57BL/6 mice. Linear regression was used to correlate C_max_ as a function of dose for plasma (blue) and ELF (red) as: Y = 0.01520*X + 3.716, and Y = 0.004664*X + 0.2826 with R^2^ = 0.6916, and 0.6667, respectively.

**Fig. 4.**
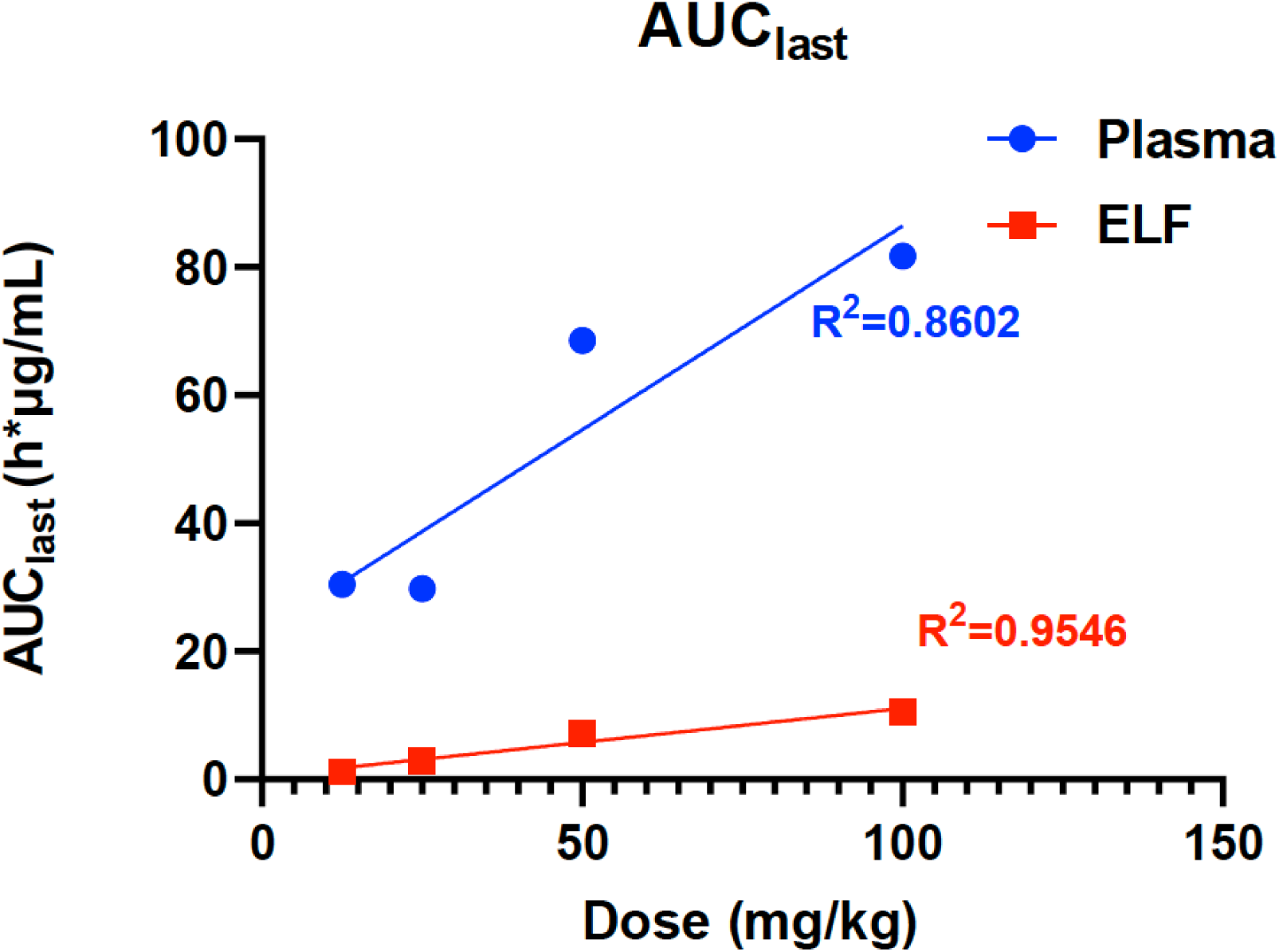
Dose correlation of ISFP10 at IP administration with AUC_last_ in plasma and ELF (µg/mL) of uninfected C57BL/6 mice. Linear regression was used to correlate AUC_last_ as a function of dose for plasma (blue) and ELF (red) as: Y = 0.6359*X + 22.82, and Y = 0.1061*X + 0.4890 with R^2^ = 0.8602, and 0.9546, respectively.

## 3. Conclusions

For the MTD of ISFP10, the tolerability of ISFP10 was evaluated in immunocompetent, uninfected male and female C57BL/6 mice. Animals received intraperitoneal (IP) administration of ISFP10 at doses of 12.5, 25, 50, 100, or 200 mg/kg, administered twice daily (BID) every 12 hours for seven consecutive days. Mortality occurred in both sexes at the 200 mg/kg dose, indicating that this concentration was not well tolerated. In contrast, doses of 12.5, 25, 50, and 100 mg/kg were tolerated by mice of both sexes without observed lethality.

Finally, this study presents the pharmacokinetic evaluation of ISFP10 in immunocompetent, uninfected C57BL/6 mice. Mice received intraperitoneal (IP) doses of ISFP10 at 12.5, 25, 50, and 100 mg/kg twice daily for seven consecutive days. ISFP10 exhibited T_max_ values ranging from 2 to 8 hours. Estimated plasma elimination half-lives (t₁/₂) were 4.39 hours and 5.31 hours for the 12.5 and 50 mg/kg groups, respectively. Plasma AUC_last_ values increased with dose, ranging from 30.463 to 81.717 h·µg/mL, indicating moderate linearity (R² = 0.8602).

In epithelial lining fluid (ELF), AUC_last_ values ranged from 1.197 to 10.561 h·µg/mL, corresponding to ELF penetration ratios (AUC_last_ ELF / AUC_last_ plasma) of 3.93% to 12.92% across the tested dosing regimens.

## 4. Conclusions

These findings, which establish the maximum tolerated dose (MTD) and pharmacokinetic (PK) profile of ISFP10 in mice, provides essential data for designing future therapeutic studies in mouse models of fungal infection. This information will guide appropriate dosing strategies, improve the interpretation of ISFP10 exposure in relation to therapeutic outcomes, and support translational efforts toward the possible clinical development of ISFP10.

## Ethics approval

These studies were approved by the Institutional Animal Care and Use Committee (IACUC) approved protocol CNS002-02232022 (MTD study) and (PK001-08062024) (PK study).

## Consent to participate

Not applicable

## Consent for publication

Not applicable

## Availability of data and material

The datasets generated and/or analyzed during the current study are available from the corresponding author on reasonable request.

## Code availability

Not applicable

## Author contributions

T.J.K. and A.H.L. made equal contributions to the conception or design of the work, and the writing of the final manuscript. Both authors approved the final version to be published.

